# QuickXL: An Open-source Ultraviolet Irradiation Device with Programmable Exposure and Active Cooling for Nucleic Acid Crosslinking

**DOI:** 10.64898/2026.11.11.766032

**Authors:** Ethan A. Kimmett, Victoria J. DeRose

## Abstract

Accessible and reproducible ultraviolet (UV) exposure is important for many chemical and biochemical applications. We describe an open-source laboratory irradiation device designed for psoralen-mediated RNA photocrosslinking, a valuable tool for nucleic acid structure determination. The device is based on 24 nominally 365–370 nm light-emitting diode modules arranged as two selectable 12-emitter series arrays with separate drivers. A Nano-compatible microcontroller, four pushbuttons and an OLED display provide user-programmable light-on and light-off intervals and an elapsed-time limit. The arrays share a fan-cooled aluminum heatsink and sample chamber, and the device can operate inside a 4 °C refrigerator to limit sample warming. A 3D-printed enclosure incorporates a sliding cover and interchangeable tube and microplate configurations with an approximately 10 mm emitter-to-sample spacing. The design supports inexpensive, configurable irradiation for nucleic-acid sample preparation, and can be adapted for other light-triggered activation of biological samples.

## Specifications table

| Hardware name | QuickXL UV Crosslinker |
|---|---|
| Subject area | • Chemistry and biochemistry |
| Hardware type | • Biological sample handling and preparation |
| Closest commercial analog | Stratagene Stratalinker UV Crosslinker 2400, AlphaThera LEDPX2 |
| Open source license | Creative Commons Attribution 4.0 International License |
| Cost of hardware | ~\$100 USD |
| Source file repository | http://doi.org/10.5281/zenodo.23653085 |

## 1. Hardware in context

Photochemical crosslinking is widely used in molecular biology to stabilize molecular interactions through the formation of covalent bonds following ultraviolet (UV) irradiation. In nucleic acid research, photocrosslinking has been particularly valuable for investigating DNA and RNA structure, RNA–RNA interactions, and nucleic acid–protein complexes [1–10]. While direct UV irradiation can induce crosslinking between nucleic acids and associated proteins, photoactivatable chemical crosslinkers provide additional selectivity for particular molecular interactions. Among these reagents, psoralens and their derivatives have been extensively employed to capture base-paired regions of DNA and RNA, enabling experimental investigation of nucleic acid secondary structure and long-range interactions [1].

Psoralens are planar, tricyclic furocoumarin compounds capable of intercalating between adjacent base pairs in double-stranded nucleic acids. Upon irradiation with long-wavelength UVA light, typically near 365 nm, psoralens undergo photochemical ‘2+2’ cycloaddition reactions with pyrimidine nucleobases, particularly thymine in DNA and uracil in RNA. Initial photoactivation produces a covalent monoadduct, while subsequent irradiation can facilitate formation of an interstrand crosslink between appropriately positioned pyrimidines on opposing strands [1]. Because these reactions require favorable nucleobase positioning within double-helical regions, psoralen photocrosslinking preferentially captures accessible nucleic acid duplexes rather than nonspecific molecular proximity. Several psoralen derivatives have been developed to improve aqueous solubility, nucleic acid affinity, and photochemical reactivity. Among these, 4′-aminomethyl-4,5′,8-trimethylpsoralen (AMT, also referred to as 4′-aminomethyltrioxsalen) is widely used for RNA structural analysis due to its improved solubility and affinity for nucleic acids. Psoralen-crosslinked RNA duplexes can be distinguished through altered electrophoretic mobility, and many psoralen crosslinks can subsequently be reversed using shorter-wavelength UV irradiation, commonly at 254 nm, to facilitate downstream analysis [2–4].

The application of psoralen photocrosslinking to RNA structural biology predates modern sequencing technologies. Calvet and Pederson demonstrated that AMT-mediated photocrosslinking could capture double-stranded regions of heterogeneous nuclear RNA in living HeLa cells, providing early evidence for structured RNA regions within native ribonucleoprotein complexes [2]. Subsequently, Thompson and Hearst used psoralen crosslinking to investigate the secondary structure of *Escherichia coli* 16S ribosomal RNA, identifying crosslinks that confirmed predicted structural features and distinguished between alternative folding models [3]. More recently, advances in high-throughput sequencing have enabled psoralen-mediated crosslinking to characterize complex RNA structures and interactions on a transcriptome-wide scale. Methods including Psoralen Analysis of RNA Interactions and Structures (PARIS) and Sequencing of Psoralen-crosslinked, Ligated, and Selected Hybrids (SPLASH) combine in-cell RNA photocrosslinking with biochemical enrichment, proximity ligation, and sequencing to identify intra- and intermolecular RNA interactions [5,6]. Crosslinking of Matched RNAs and Deep Sequencing (COMRADES) has similarly enabled investigation of alternative RNA conformations and long-range interactions within viral RNA genomes [7]. In particular, COMRADES analysis of SARS-CoV-2 revealed extensive short- and long-range RNA interactions, demonstrating the utility of photocrosslinking for understanding the structural organization of large viral genomes [8].

Despite these advances, psoralen-mediated RNA crosslinking remains subject to several practical limitations. Crosslink formation depends on nucleic acid sequence, conformation, reagent accessibility, and irradiation conditions, while the recovery of crosslinked material can be limited by inefficient photochemistry and losses during downstream sample processing. Zhang and colleagues demonstrated that improvements in psoralen chemistry and biochemical processing substantially increased the efficiency of RNA duplex detection in the PARIS2 method [9]. These findings emphasize the importance of optimizing photocrosslinking workflows to maximize recoverable crosslinked RNA while preserving sample integrity. Furthermore, prolonged UV irradiation can introduce photochemical damage and sample heating, potentially altering the experimental conditions under which RNA structures are captured [1,10]. For temperature-sensitive biological samples, the ability to maintain consistent irradiation conditions while limiting thermal accumulation is therefore an important consideration.

The effectiveness and reproducibility of photocrosslinking also depend on the characteristics of the UV irradiation system. The radiant exposure delivered to a sample is determined by the incident irradiance and illumination duration, while actual photochemical exposure additionally depends on the emission spectrum, sample geometry, vessel transmission, and spatial uniformity of illumination. Conventional laboratory UV sources, including mercury-lamp crosslinkers and handheld irradiation devices, may provide limited flexibility in source positioning, sample format, or temporal exposure control. UV light-emitting diodes (LEDs) offer an alternative that enables compact source arrangements, rapid electronic switching, and programmable irradiation schedules. Olson and colleagues recently described an open-source 365 nm LED photocrosslinking chamber designed for bioconjugation reactions in standard microcentrifuge tubes, demonstrating the feasibility of inexpensive, application-specific UV instrumentation [11]. However, nucleic acid structural studies may require different sample geometries, simultaneous treatment of many samples, and management of sample temperature during extended irradiation.

Here, we present an open-source UVA irradiation device developed for laboratory-scale nucleic acid photocrosslinking, with particular emphasis on psoralen-mediated RNA crosslinking. The system incorporates 24 nominally 365–370 nm LEDs arranged into two independently selectable arrays of 12, each powered by a dedicated constant-current driver. An Arduino-compatible microcontroller and OLED interface enable programmable control over irradiation duration, dark intervals, and total treatment time. A custom 3D-printed enclosure accommodates multiple sample formats, including standard multiwell plates, while maintaining a defined 10 mm LED-to-sample distance when used with a standard 96-well plate. The LED arrays are mounted on a shared aluminum heatsink with continuous fan cooling. For temperature-sensitive RNA applications, the entire device can be operated within a refrigerated environment, allowing heat generated during irradiation to be dissipated without an integrated active cooling system. Programmable irradiation intervals additionally enable experimental evaluation of the effects of interrupted UV exposure and thermal accumulation on crosslinking performance. Together, these features provide an accessible and adaptable platform for reproducible nucleic acid photocrosslinking and optimization of RNA structural analysis workflows.

## 2. Hardware description

QuickXL is a compact, low-cost, open-source ultraviolet A (UVA) irradiation system developed for controlled photocrosslinking of nucleic acids and related laboratory photochemical applications. The instrument combines a 24-LED UVA illumination system, programmable microcontroller-based exposure control, interchangeable sample holders, and continuous forced-air cooling within a custom 3D-printed enclosure. A photographic overview of the device is shown in Figure 1. QuickXL was developed primarily for psoralen-mediated RNA photocrosslinking, with an emphasis on reproducible illumination, flexible sample accommodation, and temperature stability during prolonged irradiation. The instrument can be operated on a laboratory benchtop or within a standard laboratory refrigerator.

**Figure 1.**
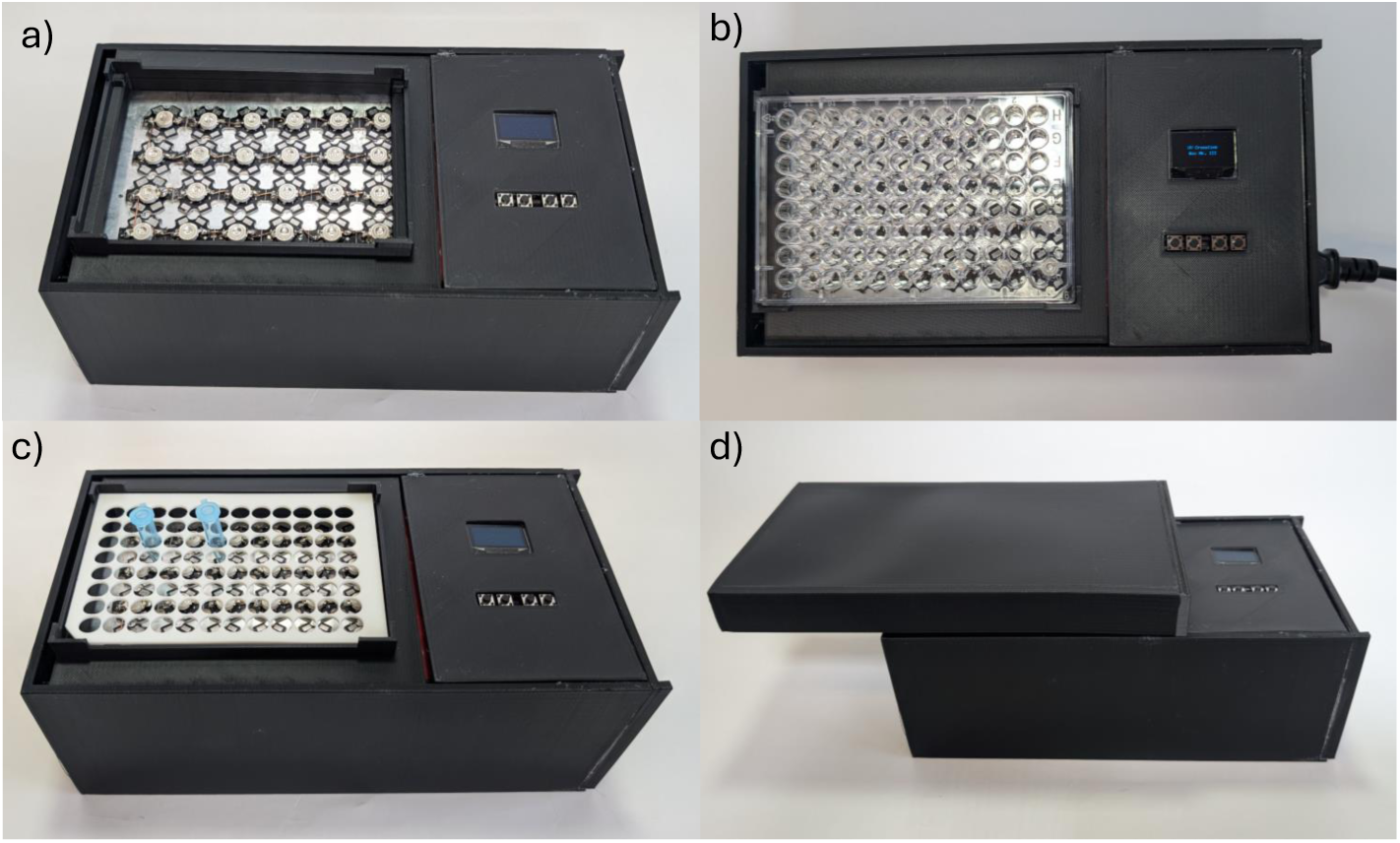
Four views of the LED field and controls, tube-support insert, microplate placement and sliding enclosure. (a) top-down view with sample holder removed, (b) top-down view 96-well plate in place, (c) top-down view with tube holder insert (d) front view with sliding cover in place in ‘half-closed’ position.

**Figure 2.**
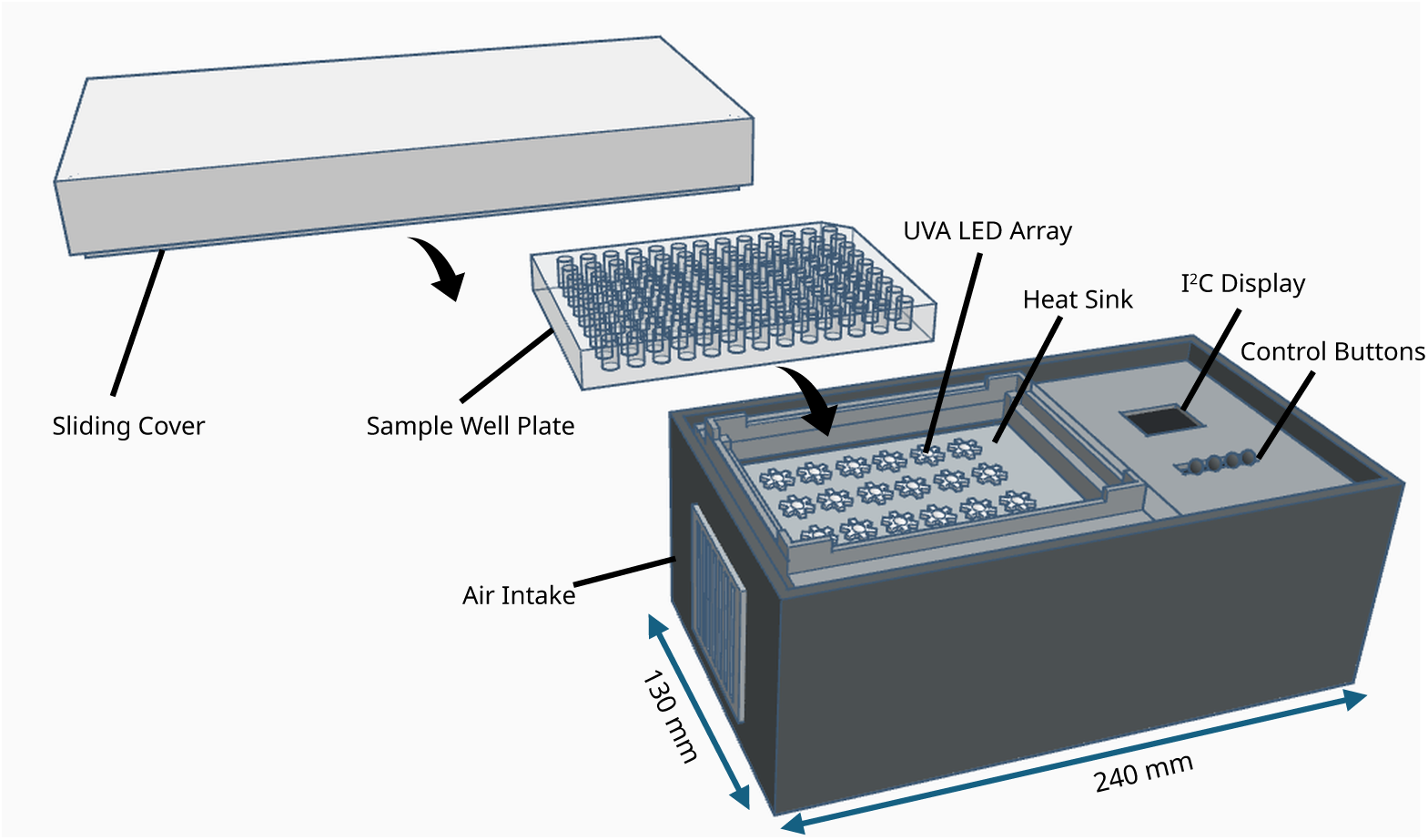
Schematic of assembled device enclosure in exploded view. Components are displaced for display.

The assembled enclosure measures 240 × 130 × 118 mm (length × width × height) and is fabricated entirely from PLA Pro, requiring approximately 620 g of filament for all 3D-printed components. The device accommodates standard 96-well plates and, through an interchangeable insert, PCR tubes and 0.6, 1.5, or 2 mL microcentrifuge tubes. Its illumination system consists of 24 nominal 3 W UVA LEDs specified as 365–370 nm emitters, with a measured peak emission wavelength of approximately 371.3 nm. The LEDs are divided into two independently selectable arrays, allowing users to control the illuminated region according to experimental requirements. Programmable exposure settings include illumination duration, intervening dark periods, and total treatment time.

The system incorporates a fan-cooled aluminum heatsink to limit thermal accumulation during irradiation. For temperature-sensitive experiments, the complete device can be placed inside a refrigerator maintained at approximately 4 °C. In a thermal characterization test, a 50 µL water sample remained at approximately 4.2 °C during 10 min of continuous irradiation with both LED arrays active. Under room-temperature conditions, a corresponding sample increased from 20.4 to 20.7 °C over the same period. These results demonstrate effective thermal management under the tested conditions without requiring integrated refrigeration or thermoelectric cooling hardwareQuickXL is assembled using commercially available electronic components, conventional wiring, and 3D-printed structural parts, without requiring a custom printed circuit board or additional machining of parts. Its modular design allows the entire LED and heatsink assembly to be removed and replaced with alternative configurations, enabling adaptation to different wavelengths, illumination geometries, or experimental requirements. The combination of programmable operation, interchangeable sample holders, and replaceable illumination modules provides an accessible and customizable platform for reproducible UVA irradiation.

Key highlights of the device include:

- Low cost, simple to assemble and operate, 3D-printed design requires no custom PCB fabrication or part machining
- Multi-modal sample stage enables use with multiwell sample plates, 0.6 – 2 mL centrifuge tubes, and PCR tube strips
- Active forced-air cooling of LED array and sample chamber provides consistent reaction temperature in refrigerated environments or at room temperature.
- Modular design allows for easy customization and fast swapping of LED array geometry and wavelength
- Programmable irradiation pulse and dark cycles enable precise control over reaction conditions

## 3. Design files summary

| Design file name | File type | Open source license | Location of the file |
|---|---|---|---|
| Air_Grates | STEP, STL | CC BY 4.0 | http://doi.org/10.5281/zenodo.23653085 |
| Display_Panel | STEP, STL | CC BY 4.0 | http://doi.org/10.5281/zenodo.23653085 |
| Enclosure_Main | STEP, STL | CC BY 4.0 | http://doi.org/10.5281/zenodo.23653085 |
| Enclosure_Sidepanel | STEP, STL | CC BY 4.0 | http://doi.org/10.5281/zenodo.23653085 |
| Holder_Support_Arches | STEP, STL | CC BY 4.0 | http://doi.org/10.5281/zenodo.23653085 |
| Lid_Main | STEP, STL | CC BY 4.0 | http://doi.org/10.5281/zenodo.23653085 |
| Lid_Sidepanel | STEP, STL | CC BY 4.0 | http://doi.org/10.5281/zenodo.23653085 |
| Sample_Holder | STEP, STL | CC BY 4.0 | http://doi.org/10.5281/zenodo.23653085 |
| Tube_Holder_Insert | STEP, STL | CC BY 4.0 | http://doi.org/10.5281/zenodo.23653085 |
| QuickXL_UI_Firmware.ino | Arduino .ino firmware | CC BY 4.0 | http://doi.org/10.5281/zenodo.23653085 |
| QuickXL_Wiring_Schematic.pdf | PDF | CC BY 4.0 | http://doi.org/10.5281/zenodo.23653085 |
| QuickXL_BOM.csv | CSV | CC BY 4.0 | http://doi.org/10.5281/zenodo.23653085 |
| Validation_Data | CSV, TIF, TXT | CC BY 4.0 | http://doi.org/10.5281/zenodo.23653085 |

- Air_Grates: Two 3-D printed protective air grates, these grates cover the openings in the main enclosure body and sidewall to protect fan and interior components while enabling airflow
- Display_Panel: 3-D printed panel for mounting of I^2^C display and push-button array
- Enclosure_Main: 3-D printed primary component of device enclosure, houses electrical components and supports heatsink and LED array.
- Enclosure_Sidepanel: 3-D printed side wall component of device enclosure, used for mounting of IEC inlet/power switch and cooling fan.
- Holder_Support_Arches: 3-D printed arches used to secure heatsink and support display panel
- Lid_Main: 3-D printed primary component of sliding lid
- Lid_Sidepanel: 3-D printed side wall component of sliding lid
- Sample_Holder: 3-D printed sample stage to support multiwell plate or tube holder insert
- Tube_Holder_Insert: 3-D printed optional insert for sample holder to support sample tubes
- QuickXl_UI_Firmware.ino: Arduino firmware for QuickXL system
- QuickXL_Wiring_Schematic.pdf: Schematic showing electrical connections in QuickXL system
- QuickXL_BOM.csv: Complete Bill of Materials for QuickXL system
- Validation Data: Raw spectral and thermal data from QuickXL characterization experiments, raw image of denaturing PAGE gel from crosslinking efficiency comparison experiment and lane assignments.

## 4. Bill of materials summary

| Designator | Component | Number | Cost per unit - currency | Total cost - currency | Source of materials | Material type |
|---|---|---|---|---|---|---|
| LED | Nominal 3 W UVA modules | 24 | 28.80 | 1.20 | https://www.amazon.com/dp/B0CSSKPVHC/ | Other |
| DRV | LED driver, as-installed model | 2 | 19.98 | 9.99 | https://www.amazon.com/dp/B01N6U6A4E/ | Other |
| RELAY | LED-driver AC-input switching module | 2 | 3.50 | 1.75 | https://www.amazon.com/dp/B095YD3732/ | Other |
| MCU | ELEGOO Nano USB-C controller | 1 | 5.33 | 5.33 | https://www.amazon.com/dp/B0F6Y7GS4Q/ | Other |
| DISP | 128 × 64 I²C SSD1306 OLED Display | 1 | 3.00 | 3.00 | https://www.amazon.com/dp/B09C5K91H7/ | Other |
| PSU | AC-to-5 V DC supply | 1 | 2.00 | 2.00 | https://www.amazon.com/dp/B0B2RBP1Z2/ | Other |
| HS | Aluminum heatsink | 1 | 10.99 | 10.99 | https://www.amazon.com/dp/B08JQGSXG3/ | Other |
| ADH | Thermal adhesive | 10g | 1.60 | 7.99 | https://www.amazon.com/dp/B0DL5X4Y7P | Other |
| INLET | IEC C14 inlet / switch / fuse | 1 | 2.10 | 2.10 | https://www.amazon.com/dp/B082ZFRV1B/ | Other |
| FAN | Continuous cooling fan | 1 | 2.25 | 2.25 | https://www.amazon.com/dp/B09Y5Q2LLG | Other |

The complete Bill of Materials (QuickXL_BOM.csv) is available at the following Zenodo repository: http://doi.org/10.5281/zenodo.23653085. All components were obtained from Amazon (US), with a total allocated cost of 96.97 USD as of October 2026. Equivalent parts are available from other electronics distributors worldwide.

## 5. Build instructions

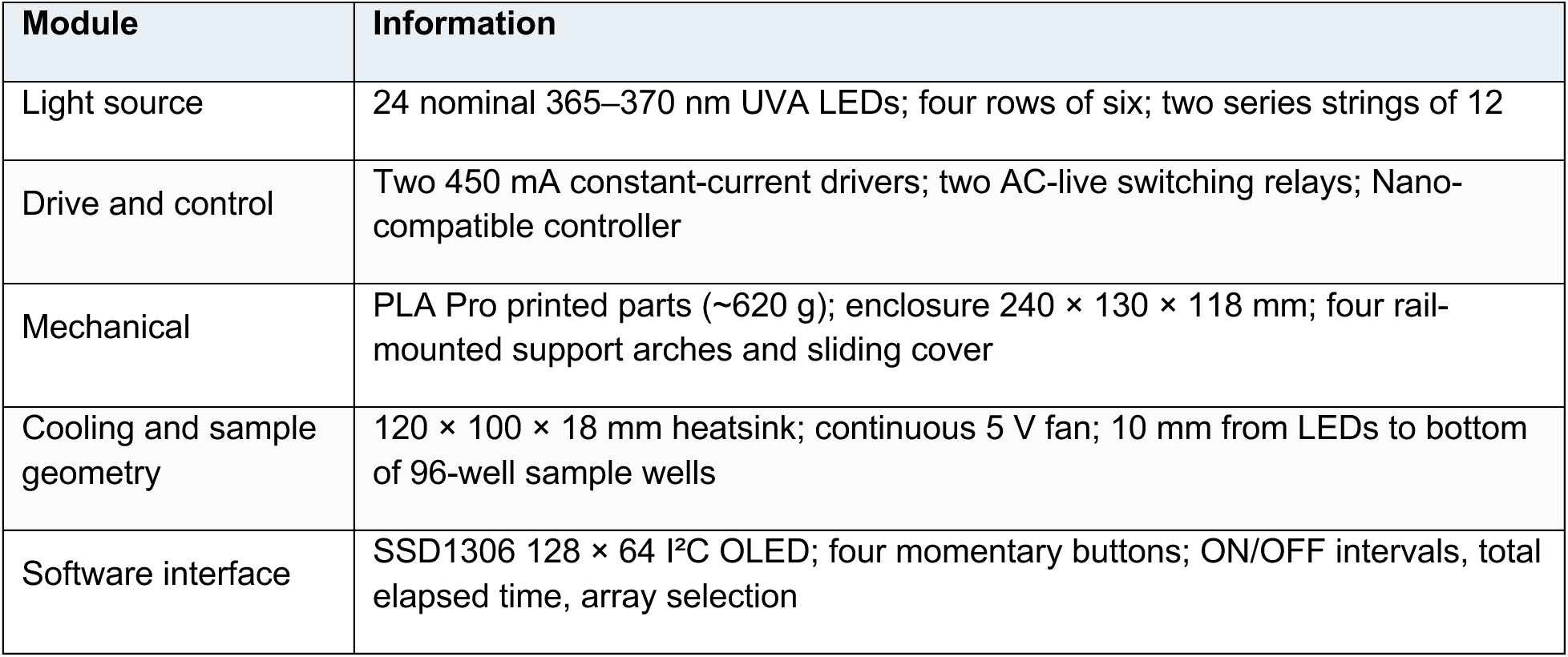

| **Module** | **Information** |
|---|---|
| Light source | 24 nominal 365–370 nm UVA LEDs; four rows of six; two series strings of 12 |
| Drive and control | Two 450 mA constant-current drivers; two AC-live switching relays; Nano-compatible controller |
| Mechanical | PLA Pro printed parts (~620 g); enclosure 240 × 130 × 118 mm; four rail-mounted support arches and sliding cover |
| Cooling and sample geometry | 120 × 100 × 18 mm heatsink; continuous 5 V fan; 10 mm from LEDs to bottom of 96-well sample wells |
| Software interface | SSD1306 128 × 64 I²C OLED; four momentary buttons; ON/OFF intervals, total elapsed time, array selection |

### Safety and prerequisites

#### 5.1 Critical precautions

- AC mains input is present inside this instrument. Build and inspect the mains side only with the device fully disconnected.
- The device emits UVA around 365–370 nm. Never view energized LEDs or run with the sample area exposed. The correct START configuration is HALF-CLOSED: the cover completely shields the emitter/sample region while leaving the display/buttons accessible. After pressing Enter to start, slide the cover to FULLY CLOSED over the display. Verify UVA containment at the half-closed and fully closed positions before use.
- During refrigerated operation, prevent condensation, drips, and liquid spills from reaching the mains or low-voltage compartments. Do not route a mains power lead through an unapproved refrigerator opening.

#### 5.2 Parts and tools

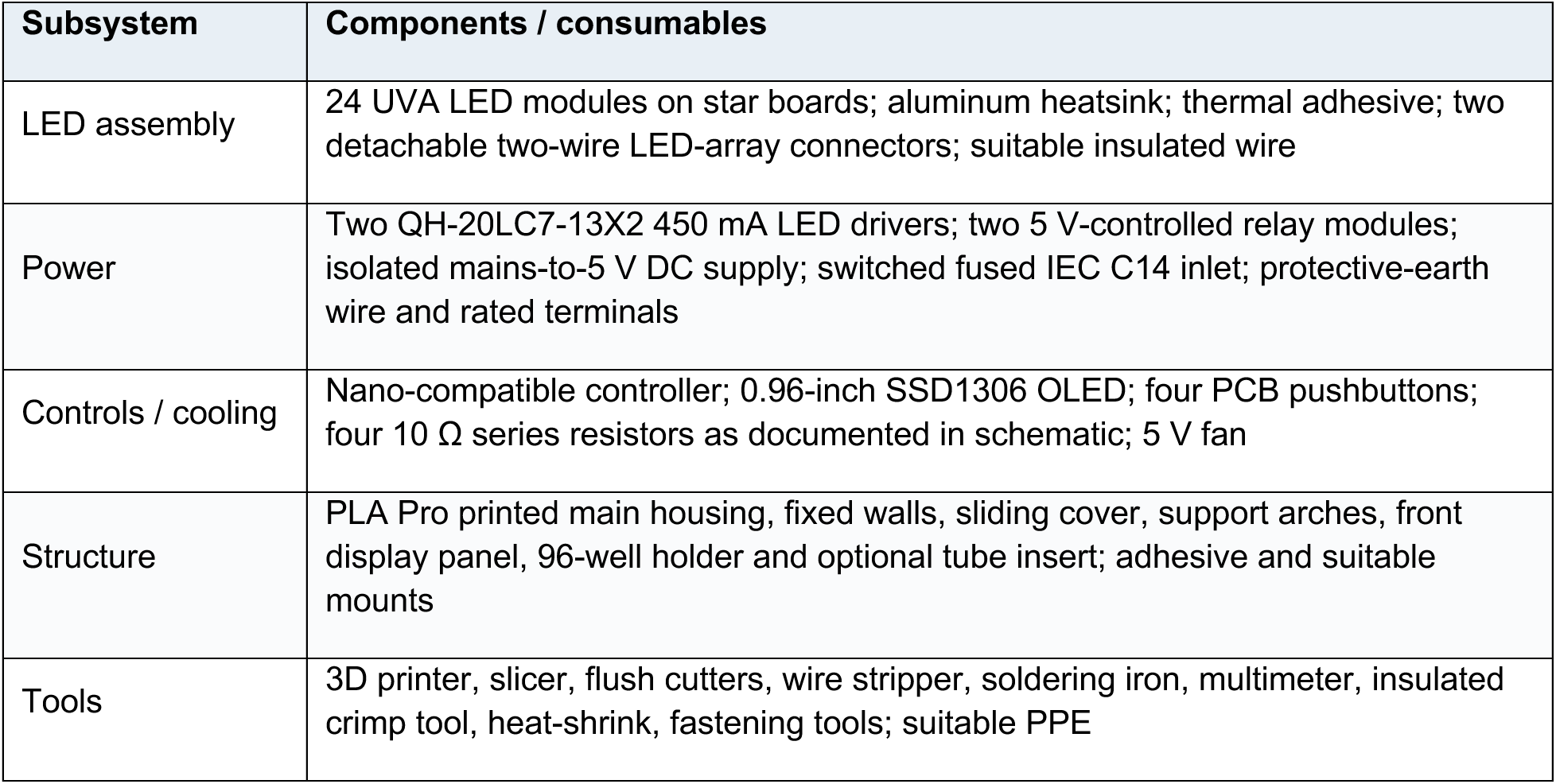

### Mechanical construction

#### 5.3 Print and identify the plastic parts

- Print from the nine named STL files supplied with this guide, keeping their millimetre geometry unchanged. Optimal settings for PLA Pro are a 0.20 mm layer height with a 0.40 mm nozzle and 25% infill, three walls, and standard supports. Verify print orientation, supports and wall count for each mesh against the slicer preview. Consult the dedicated printed-parts catalog immediately below for identification and component count.
- Print the enclosure and lid bodies, the fixed panels, the sample holder, the display panel and the optional tube insert as required. One print of Holder_Support_Arches.stl contains FOUR physically separate arches. One print of Air_Grates.stl contains TWO ventilation grilles. Confirm adequate bed size, bed adhesion and separation between the pieces in multi-part STL files.

#### 5.4 Printable component catalog

The nine files below form the supplied printed-part set. Each figure shows an isometric view beside an orthographic view. The stated X × Y × Z measurements are the file’s axis-aligned mesh extents in millimetres.

**Part P1. Main enclosure shell**

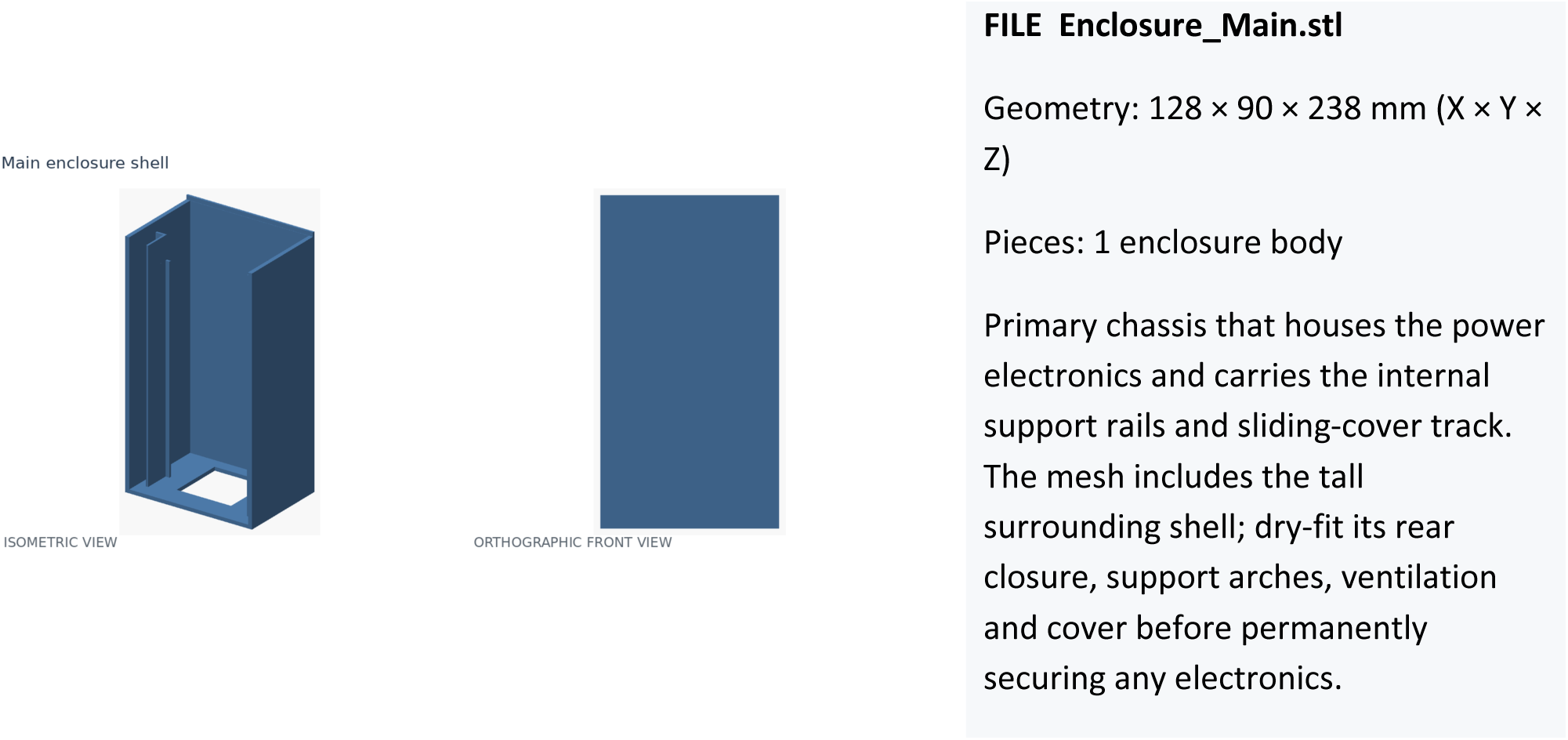

**Part P2. Fixed enclosure end/side panel**

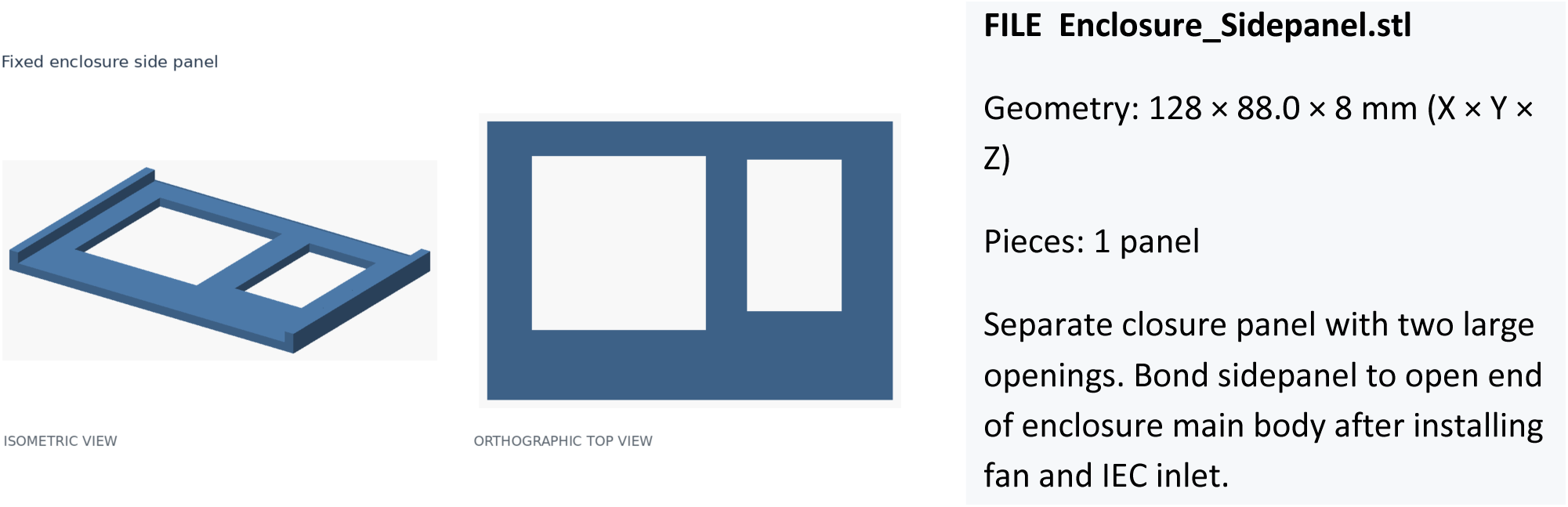

**Part P3. Sliding protective cover**

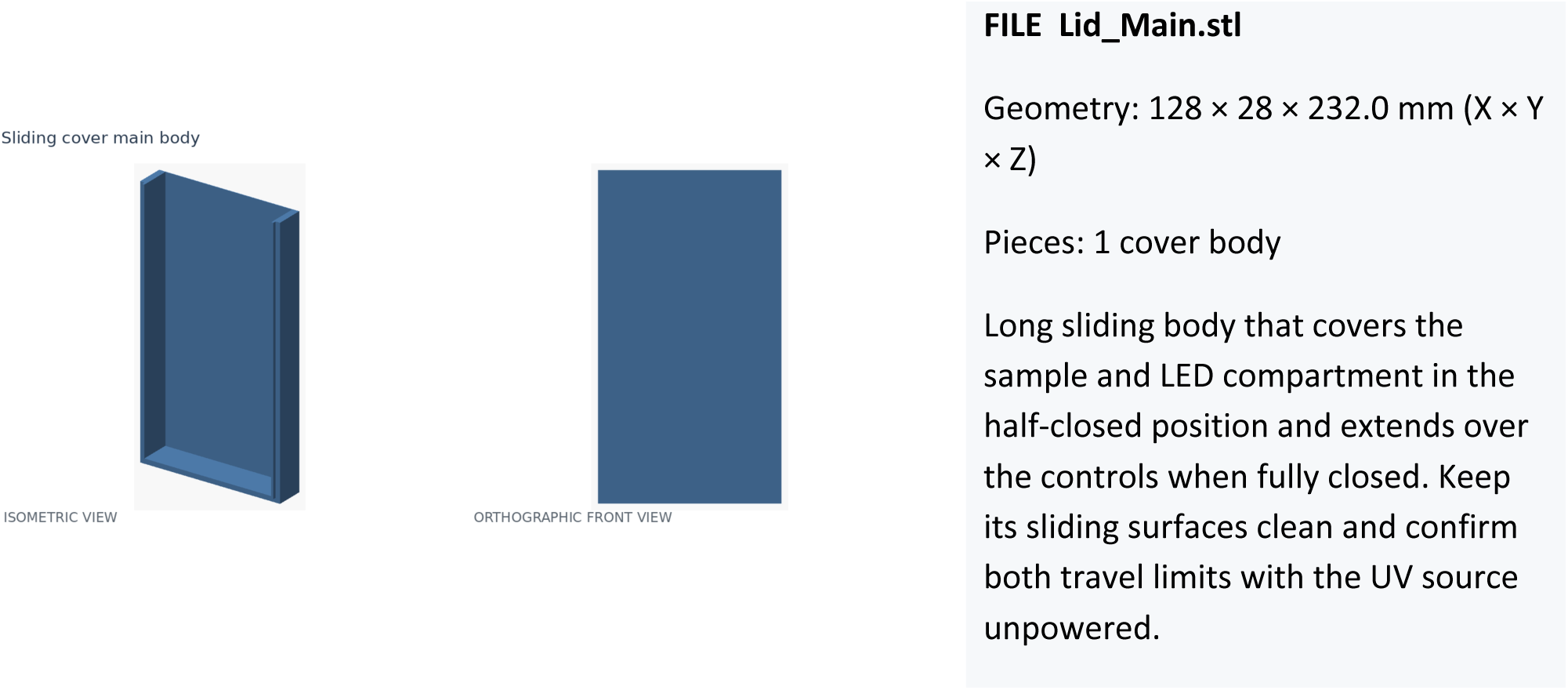

**Part P4. Sliding-cover side/endpiece**

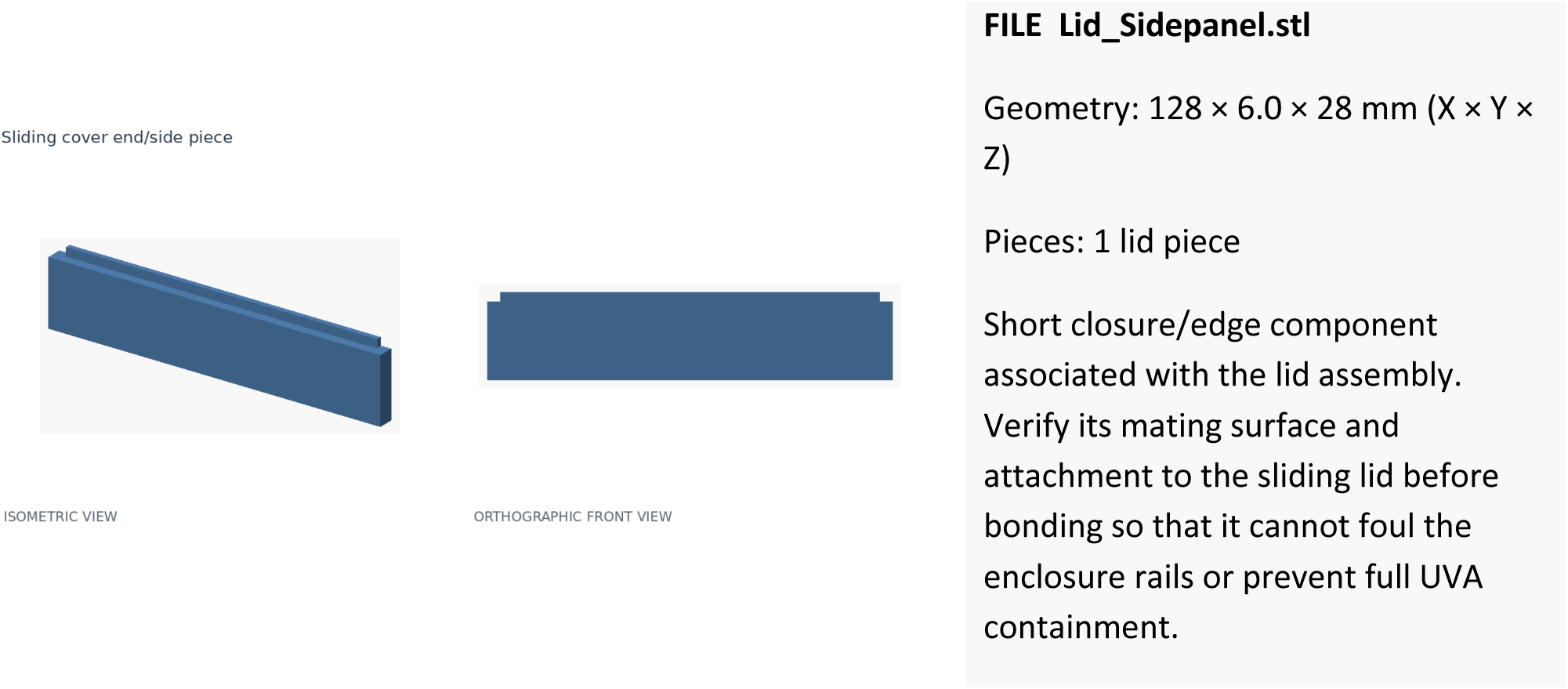

**Part P5. Four rail-mounted structural arches**

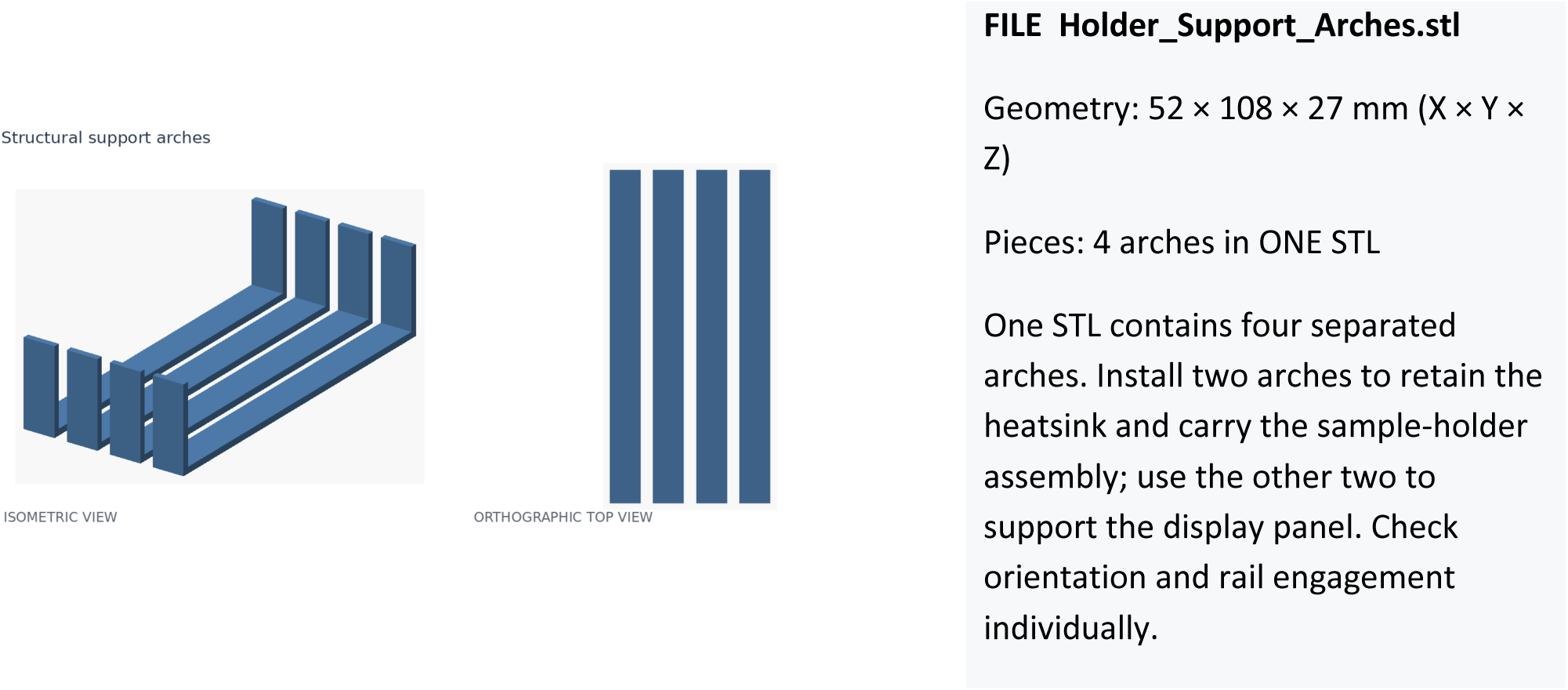

**Part P6. 96-well plate carrier**

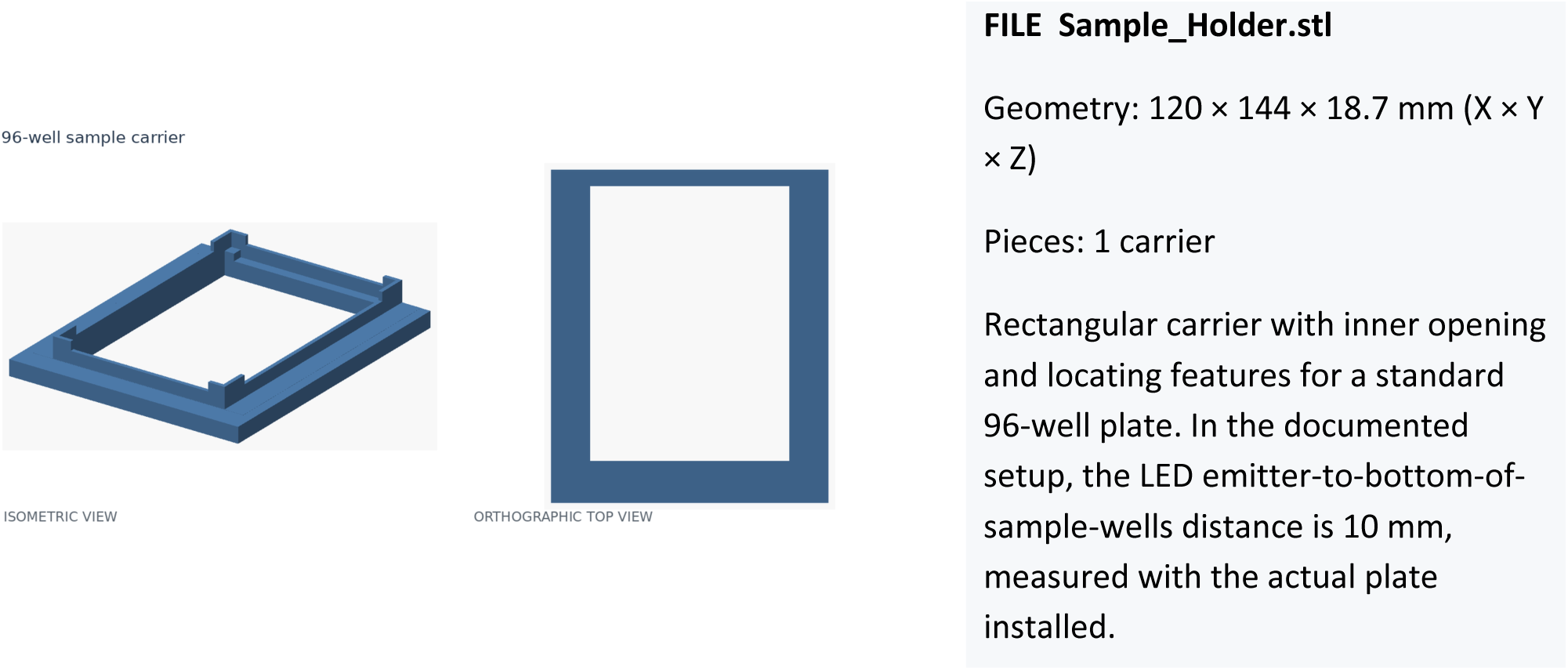

**Part P7. Optional perforated tube insert**

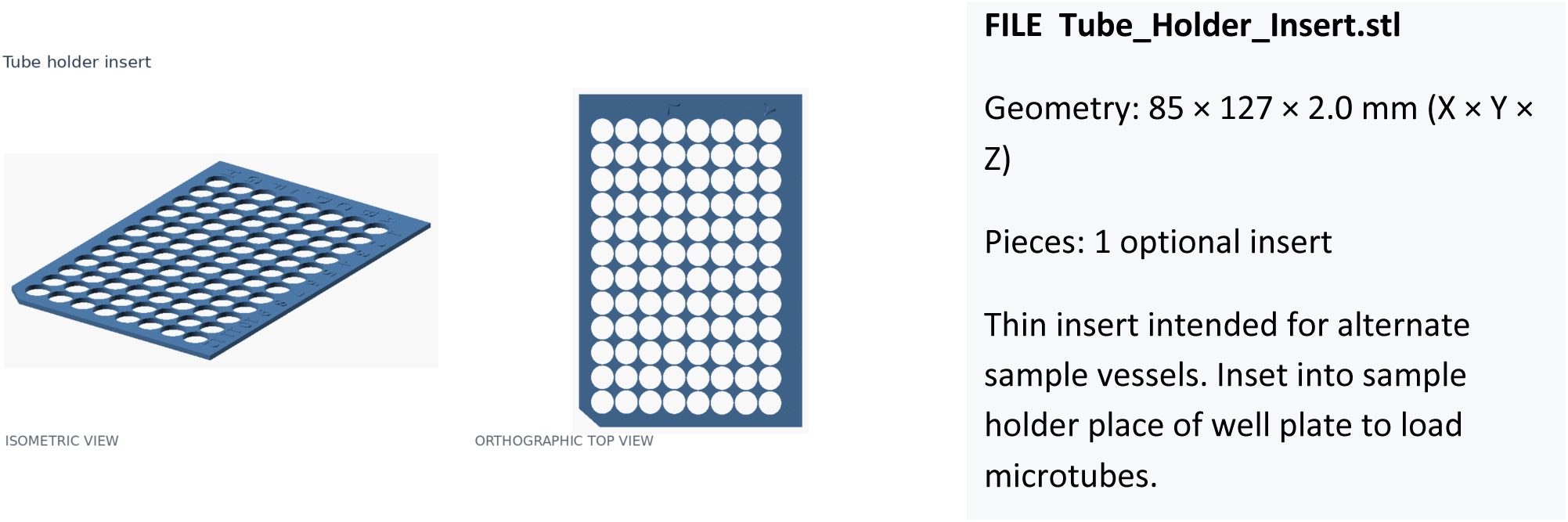

**Part P8. OLED and four-button control panel**

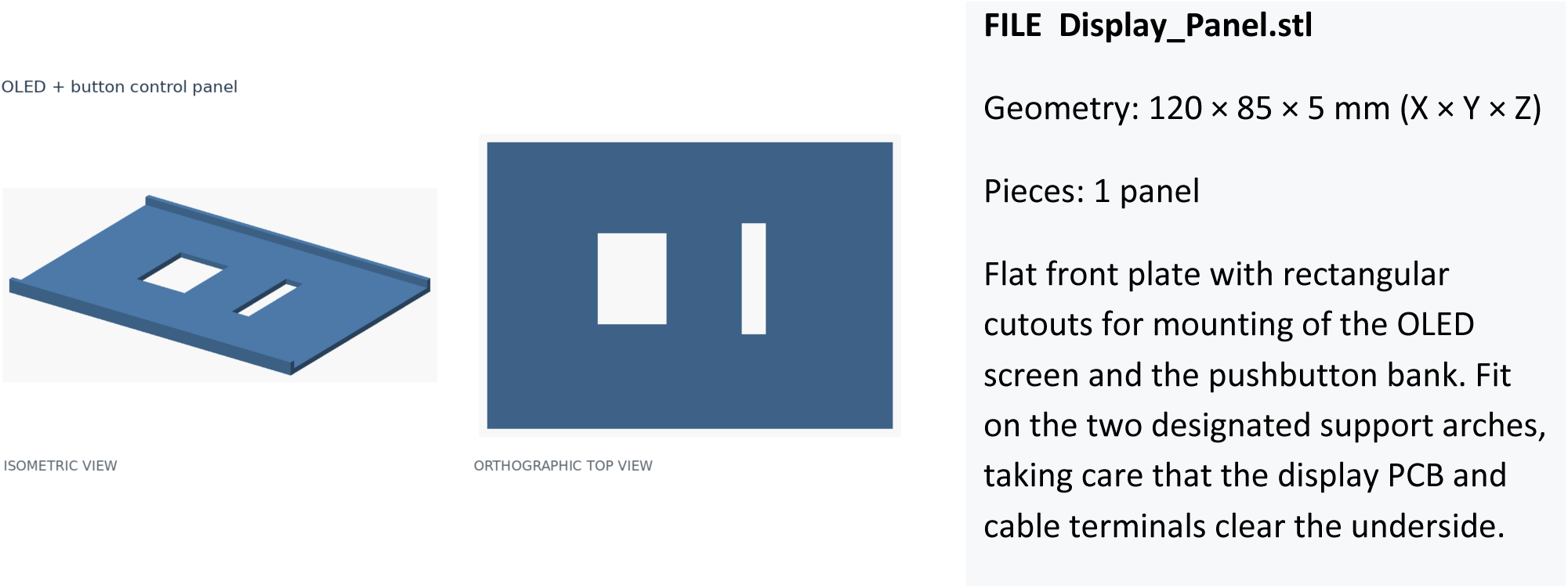

**Part P9. Two ventilation grilles**

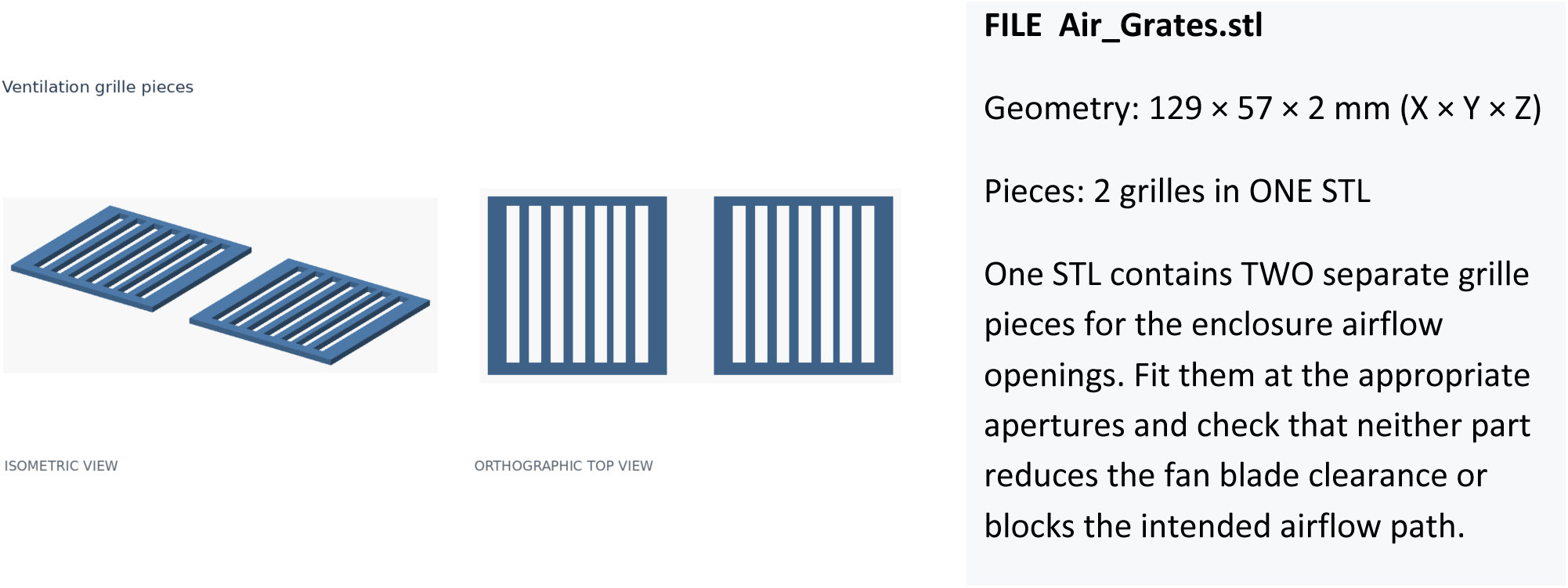

#### 5.5 Dry-fit the enclosure, inlet and sample area

- Identify permanent enclosure components (enclosure P1, side panel P2, control/display panel P8, air grates P9) and the pieces that must remain movable or removable (sliding cover P3/4, sample holder P6, support arches P5, optional tube insert P7). Check smooth operation of sliding cover P3 on main enclosure P1. Dry-fit and then bond sliding cover P3 with lid sidepanel P4. Dry-fit main enclosure P1 and side panel P2, control/display panel P8, and sample holder P6. Bond enclosure and sidepanel P1 and P2, reserving control panel P8 until after component wiring.
- Locate the control/display-panel opening (on P8), fan vent and IEC inlet cutout (on P2), heatsink rails and sample-holder rails (P1 interior). The four support arches P9 will be placed into enclosure support rails at equal intervals to separately support the heatsink/sample holder and the control/display panel. Confirm that the IEC inlet and fan can be seated without obstructing cable routing or the heatsink (Figure 5).
- Set a standard 96-well plate in its intended holder and check that the plate can be inserted without binding or rocking. Test the optional tube insert separately.

#### 5.6 Lay out the two 12-emitter LED arrays

- Place the clean 120 × 100 × 18 mm heatsink on a protected flat surface. Mark the layout of 24 star-board LEDs in four rows of six, following the photographs and released design (Figure 6).
- As confirmed for the finished device, the TWO UPPER ROWS (12 LEDs total) form the Top channel and the TWO LOWER ROWS (12 LEDs total) form the Bottom channel. Mark the independent series paths and lead exits; use matching Top/Bottom labels on the connectors, drivers and relay control leads.
- Align the modules to keep the active emitting surfaces in the intended plane; leave room for solder connections and strain relief.

#### 5.7 Bond and series-connect the LED modules

- Using the selected thermally conductive adhesive, prepare mating surfaces as directed by the product instructions; apply a uniform bond that does not cover electrical pads. Seat the LEDs and allow the adhesive to fully cure.
- Connect each channel as a series string of 12 emitters, respecting the actual anode (+) and cathode (−) markings on each board. The positive end of each completed string connects to its corresponding driver LED+ output and the negative end to LED−, through an insulated detachable connector.
- Insulate each joint, restrain the leads away from sharp heatsink edges, and label both connector halves. Check the series string with a suitable unpowered test method; do not apply AC mains during this check.

#### 5.8 Fit the cooling fan and inlet hardware

- Insert the 5 V fan into the main enclosure rear ventilation opening (next to the IEC inlet) so that airflow is directed from inside the device to the exterior (Figure 5). Secure the fan by bonding without blocking fan blades.
- Fit the switched/fused IEC C14 inlet into its designated opening and bond securely (Figure 5).
- Route the fan lead toward the 5 V distribution point. Ensure the fan spins freely by hand and all inlet/fan mounts are mechanically secure (Figure 7).

### Control/display panel and microcontroller

#### 5.9 Mount the four PCB buttons and OLED display

- Place four momentary PCB buttons into the Control panel P8 openings, maintaining the pushbutton spacing and orientation shown in the annotated photos (Figure 3 and 8). Affix them without depositing adhesive into the moving switch mechanisms.
- Mount the 128 × 64 SSD1306 OLED with its active viewing window centered in the control/display panel P8 OLED opening. Avoid applying force to the display glass, ribbon or connector pins.
- Prepare the four signal wires, button ground returns and OLED four-wire harness.
- Label the controller-side wires by function; button-to-pin relationships are defined in Figure 8.

#### 5.10 Connect the Nano-compatible controller

- Locate the controller pin labels on the particular Nano-compatible board. Bring the regulated 5 V supply to the controller 5V input and the 5 V supply return to GND as documented (Figure 9).
- Connect OLED SDA to A4 and SCL to A5, with OLED power to the appropriate 5 V rail and its ground to common GND.
- Connect the button inputs by named function: D5 = minus; D4 = plus; D3 = Back; D2 = Enter. Each button is a normally open path to GND, with the internal controller pull-up enabled in firmware.
- Reserve D8 for the Top driver relay signal and D9 for the Bottom driver relay signal. The relay logic-side supplies must share the same reference as the controller.

### Power distribution and LED switching

#### 5.11 Plan separation of AC and low-voltage wiring

- Before permanent installation, choose the power-component locations in the lower enclosure: IEC inlet/switch/fuse at the rear; two LED drivers; two relay modules; an AC-to-5 V DC converter; microcontroller and harnesses (Figure 17).
- Keep mains-voltage input and relay-contact conductors distinguishable from extra-low-voltage control, fan and OLED/button wires. Ensure appropriate insulation, secure terminals, strain relief and protected routing so the two wiring classes cannot contact each other.

#### 5.12 Install the IEC power entry and protective earth

- Verify actual inlet terminal map from the molded markings and manufacturer documentation. Identify incoming live (L), neutral (N), and protective earth (PE) (Figure 13); verify the internal switch and fuse positions electrically while the assembly is disconnected.
- The intended mains path is: fused/switched live to the individual relay contact inputs and AC-to-5 V supply as required; unswitched neutral to the driver/AC-to-5 V neutral inputs. Each relay switches only its respective LED-driver live input.
- Bond protective earth to the exposed conductive heatsink as documented. Keep PE continuous and independent of switching. Use approved mains terminals, wire sizing, protection and strain relief for the installation environment.

#### 5.13 Wire the regulated 5 V supply and continuous fan

- Identify and verify the AC-L and AC-N input terminals of the AC-to-5 V converter (Figure 14). Connect its inputs to the approved mains distribution path.
- On the extra-low-voltage side, identify regulated +5 V and DC 0 V/GND. Distribute these to the Nano, OLED, relay logic modules and fan with correct polarity.

#### 5.14 Connect the two relay control and AC switching channels

- Connect the Top relay logic input to Nano D8 and the Bottom relay logic input to Nano D9; connect each relay logic VCC to the verified +5 V supply and its logic GND to DC GND (Figure 15).
- On each relay contact side, route the approved supply live through the normally open contact path to that channel’s LED-driver live input. Keep each LED-driver neutral connected to AC-N.
- Verify the relay board terminal labels (COM, NO, NC), its actual contact configuration and any high/low-trigger selector. The firmware issues HIGH to command ON and LOW to command OFF; the installed modules must match this polarity.

#### 5.15 Connect the two constant-current LED drivers

- Identify the AC input leads of each QH-20LC7-13X2 driver and the separate DC LED+ (red) and LED− (black) outputs from the label on the installed unit (Figure 16).
- Connect each driver AC live lead to its own relay-switched AC-L; connect its input neutral to AC-N. Route the isolated DC output to the matching labeled 12-LED string through the detachable pair connector.
- Secure the drivers so AC-input and DC-output leads are not strained

#### 5.16 Perform unpowered electrical and mechanical inspection

- With AC power unplugged and energy-storage components safely discharged, verify wire labels, connector polarity, insulation and strain relief. Confirm no conductor is trapped beneath a printed panel or heatsink.
- Inspect the conductor separation, exposed metal bonding, AC-input contact covers, relay configuration and control wiring.

#### 5.17 Load and verify the current firmware

- Install the Arduino IDE and the Adafruit_GFX and Adafruit_SSD1306 libraries needed by the released firmware. Select the exact Nano-compatible board used, processor/bootloader, and USB serial port; record the IDE and library versions.
- Compile and upload the released .ino sketch (QuickXL UI Firmware.ino). Confirm that its I²C address is 0×3C and that its D2–D5, D8/D9 and active-high relay definitions match the as-built wiring.
- With LED driver mains inputs safely isolated, check OLED startup, minus/plus/Back/Enter buttons, array-selection indications, relay-off state after setup, and Back-to-STOP handling.

#### 5.18 Mount the electronics and close the fixed enclosure structure

- After successful unpowered inspection and low-voltage checks, mount the two LED drivers, two relay modules, 5 V supply and Nano into the lower Main enclosure (P1/2) in the documented arrangement (Figure 17), using appropriate epoxy adhesive
- Route the LED driver output leads toward the upper LED module mounting area, leaving sufficient slack for the detachable connectors.
- Check the enclosure interior against the overall device photograph before installing the heatsink.

#### 5.19 Install the heatsink/LED assembly and support arches

- Insert the finished LED/heatsink assembly (Figure 6) into the designated upper structure of the Main enclosure (Figure 3). The aluminum heatsink is supported by the enclosure support rails
- Fit TWO of the four support arches into the enclosure support rails at the heatsink/sample-holder location. These two arches retain the heatsink and support the sample-holder structure above it.
- Attach each LED-array connector to its labeled driver output, maintaining the previously verified channel identity and polarity. Ensure no loose cable obstructs fan blades or the illuminated field.
- Install the remaining TWO printed support arches in the rails supporting the control/display panel. Seat the wired control/display panel (Figure 8) on these arches, keeping the OLED, pushbuttons and harness accessible; check the panel is level and its leads are not pinched.
- Fit Sample holder P6 above the LED array, centered below the intended standard 96-well plate.
- Test all cover positions: OPEN for access only with UV off; HALF-CLOSED covering the entire UV/sample area but leaving the OLED/buttons exposed for setup and starting a run; FULLY CLOSED covering both UV/sample area and control panel during the run. The cover must slide without lifting or exposing the sample area during the transition.

### Controls and operation

#### 5.20 Controls

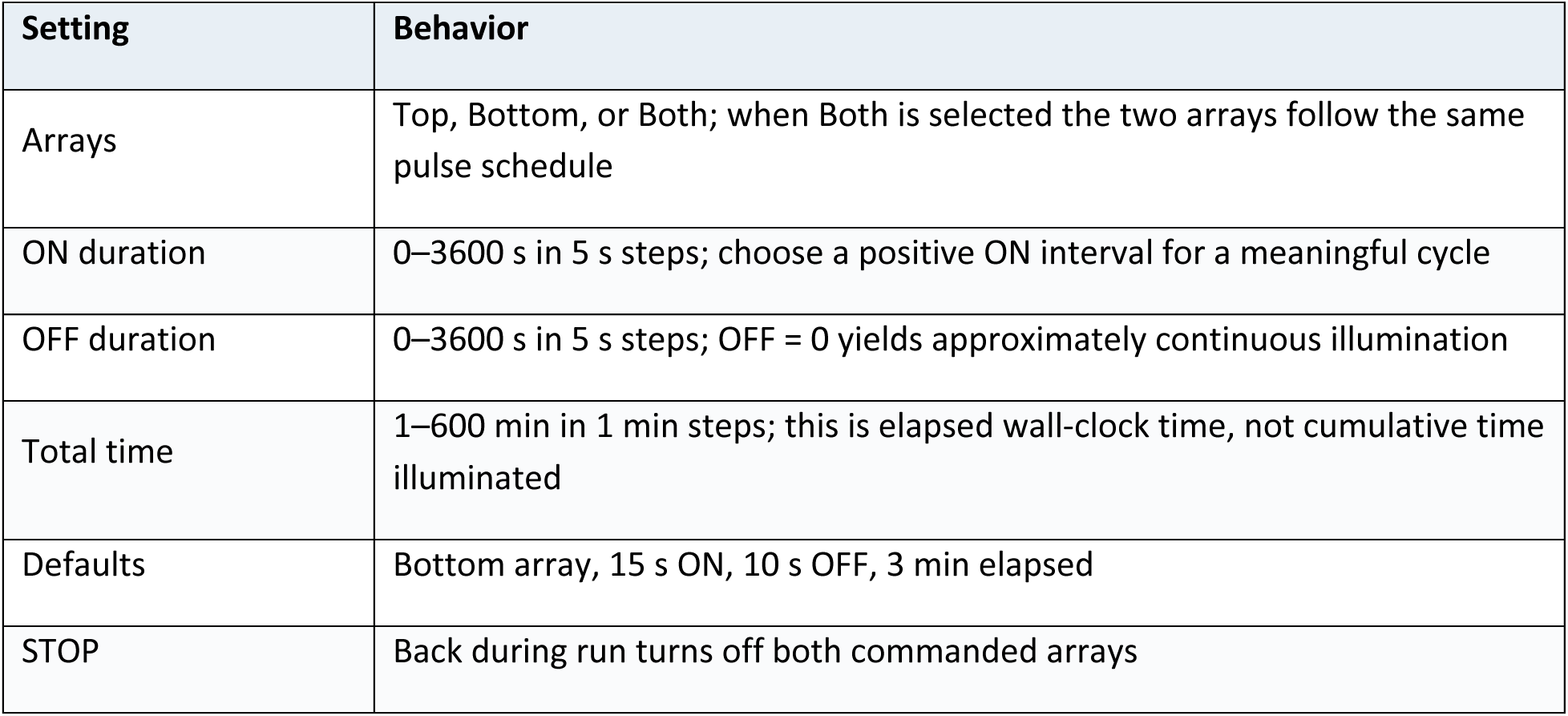

| **Setting** | **Behavior** |
|---|---|
| Arrays | Top, Bottom, or Both; when Both is selected the two arrays follow the same pulse schedule |
| ON duration | 0–3600 s in 5 s steps; choose a positive ON interval for a meaningful cycle |
| OFF duration | 0–3600 s in 5 s steps; OFF = 0 yields approximately continuous illumination |
| Total time | 1–600 min in 1 min steps; this is elapsed wall-clock time, not cumulative time illuminated |
| Defaults | Bottom array, 15 s ON, 10 s OFF, 3 min elapsed |
| STOP | Back during run turns off both commanded arrays |

**Figure 3.**
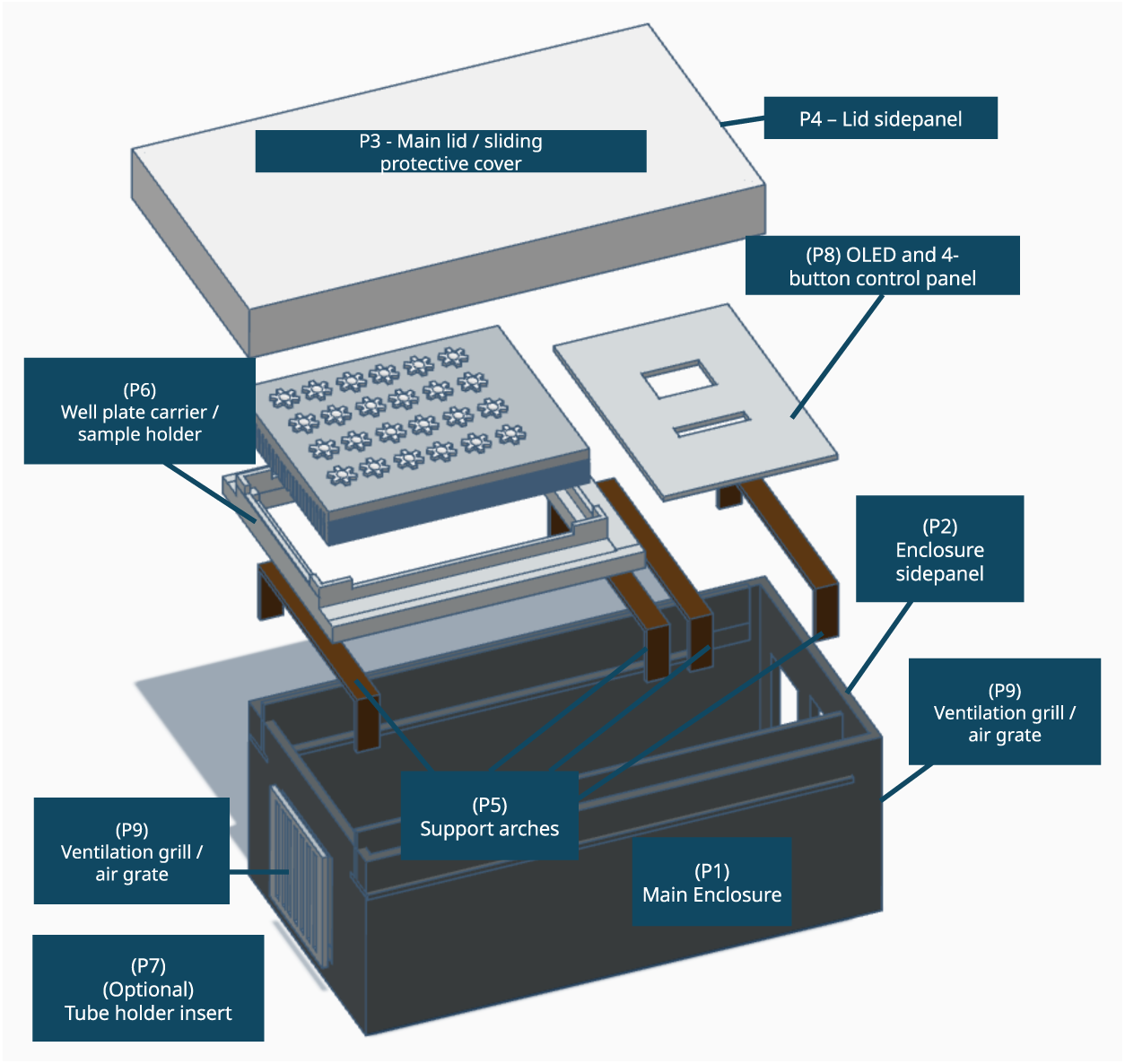
Overview of device showing assembly parts **P1-9**

**Figure 4.**
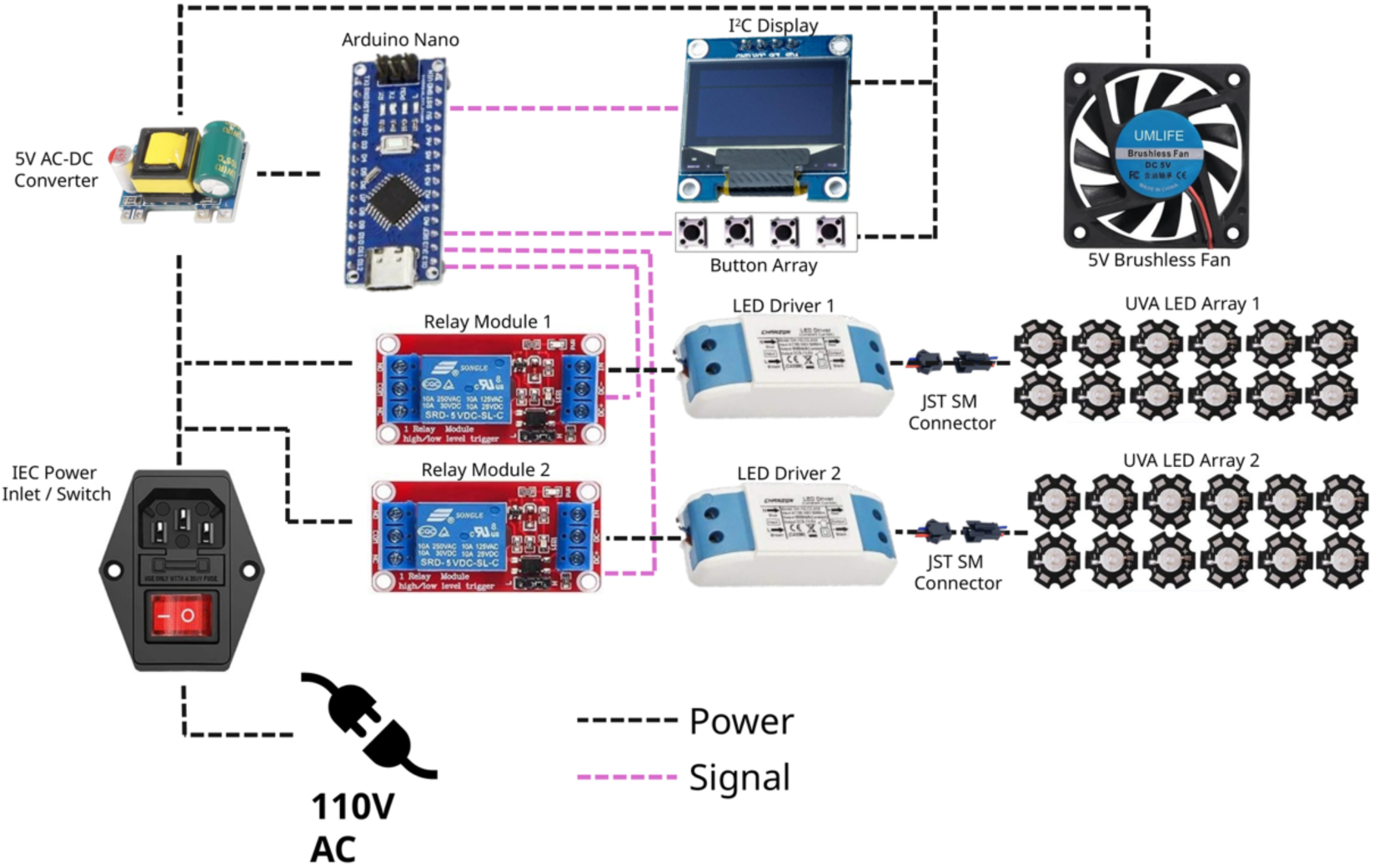
Overview of programmable device components-AC power distribution and LED driver control circuitry. Mains power enters through a fused IEC inlet and switch, supplying two independently controlled constant-current LED drivers and a 5 V AC-DC power supply. Two Arduino-controlled, normally open relays independently switch the AC live connections to the LED drivers. An Arduino Nano-compatible microcontroller controls the relays through digital outputs D8 and D9 and communicates with a 128 × 64 SSD1306 OLED display via I²C (A4/A5). Four pushbuttons connected to digital inputs D2–D5 through 10 Ω series resistors allow adjustment of irradiation parameters and operation of the device. The 5 V supply powers the microcontroller, relay modules, OLED display, and continuously operating cooling fan. Twenty-four UVA LEDs are divided into two independently controlled arrays of 12 LEDs connected in series, each powered by a dedicated constant-current driver. Both arrays can be operated individually or simultaneously using the programmed irradiation schedule.

**Figure 5.**
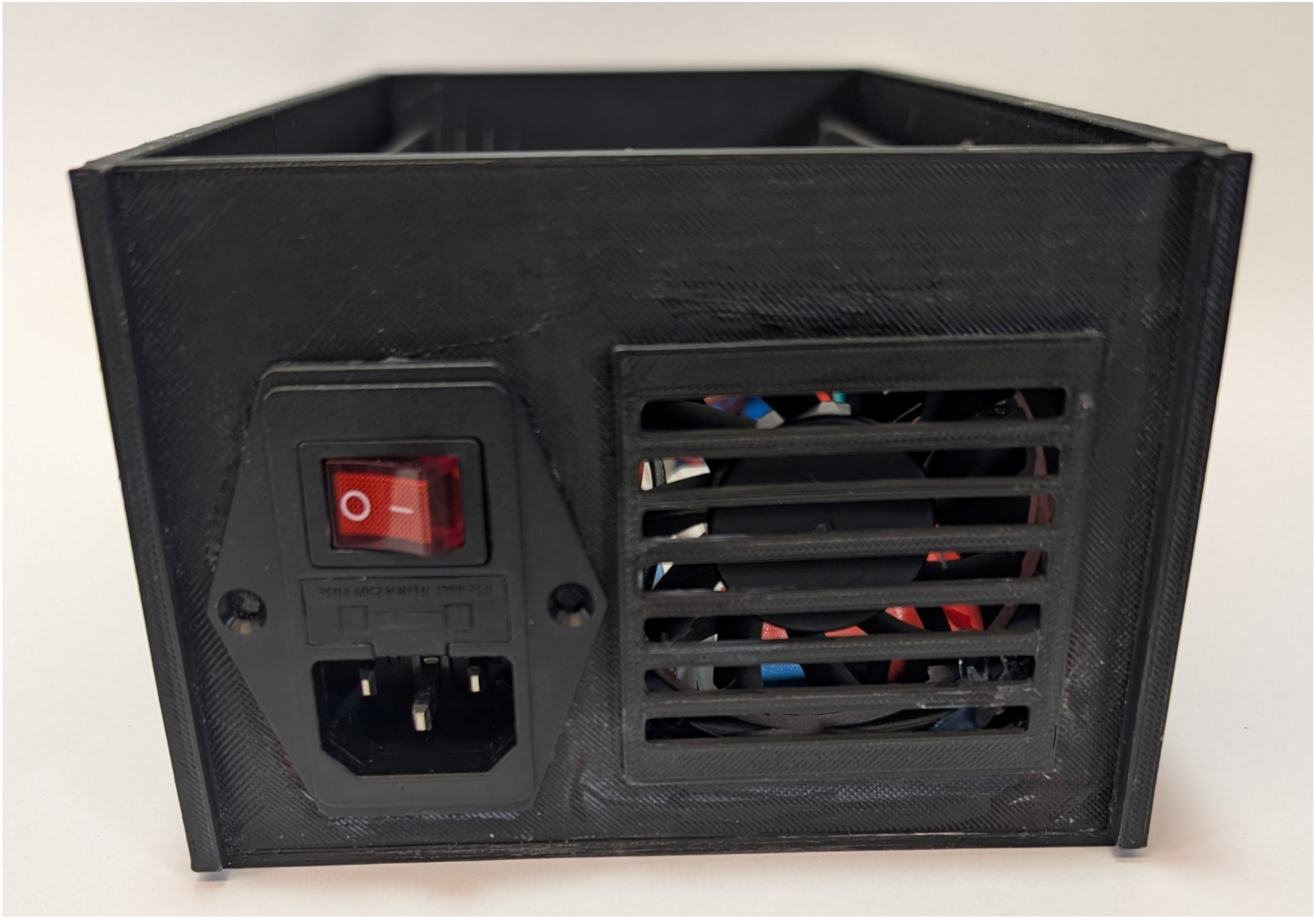
Rear view of the printed enclosure (side panel P2) with the switched IEC inlet and fan grille P9 locations.

**Figure 6.**
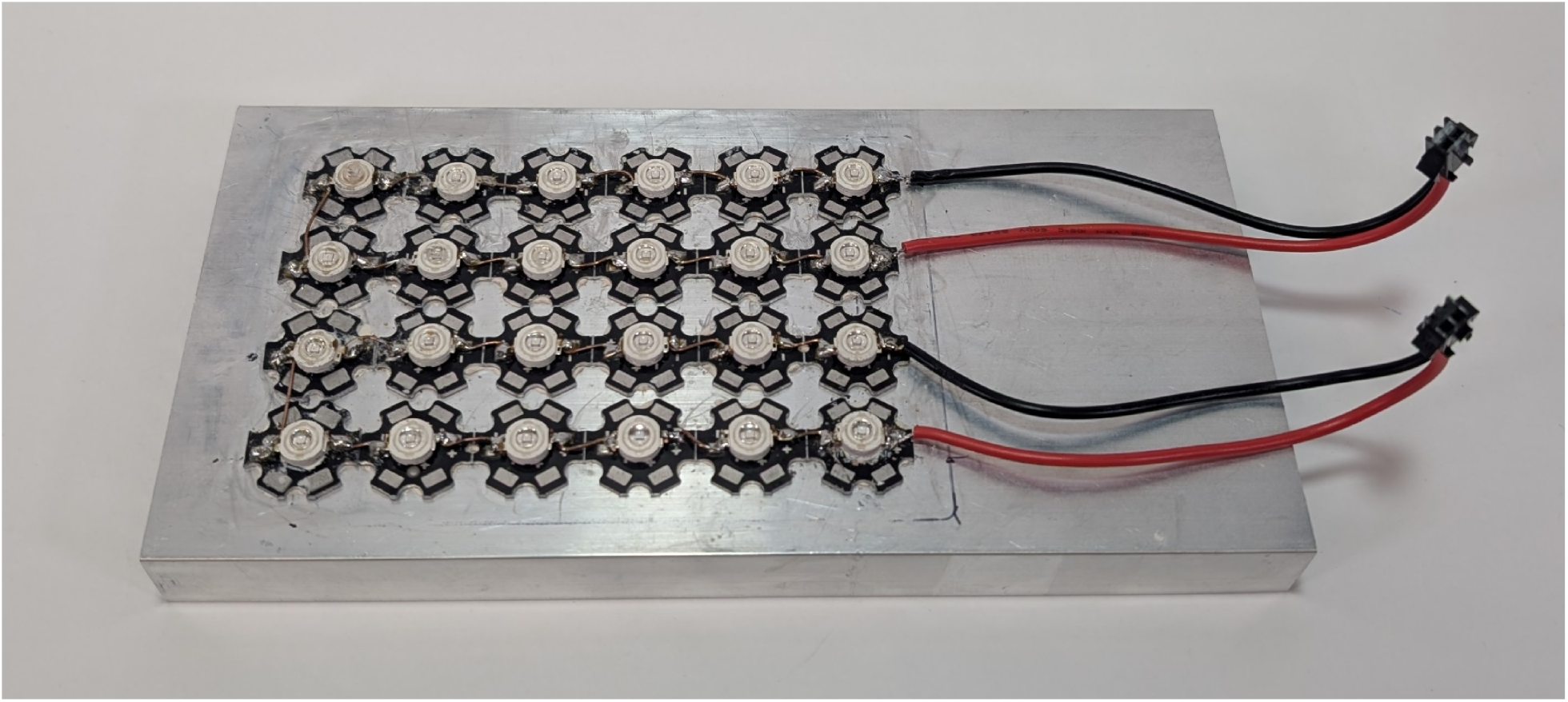
Twenty-four UVA modules: two upper rows form Top and two lower rows form Bottom, each with a dedicated connector.

**Figure 7.**
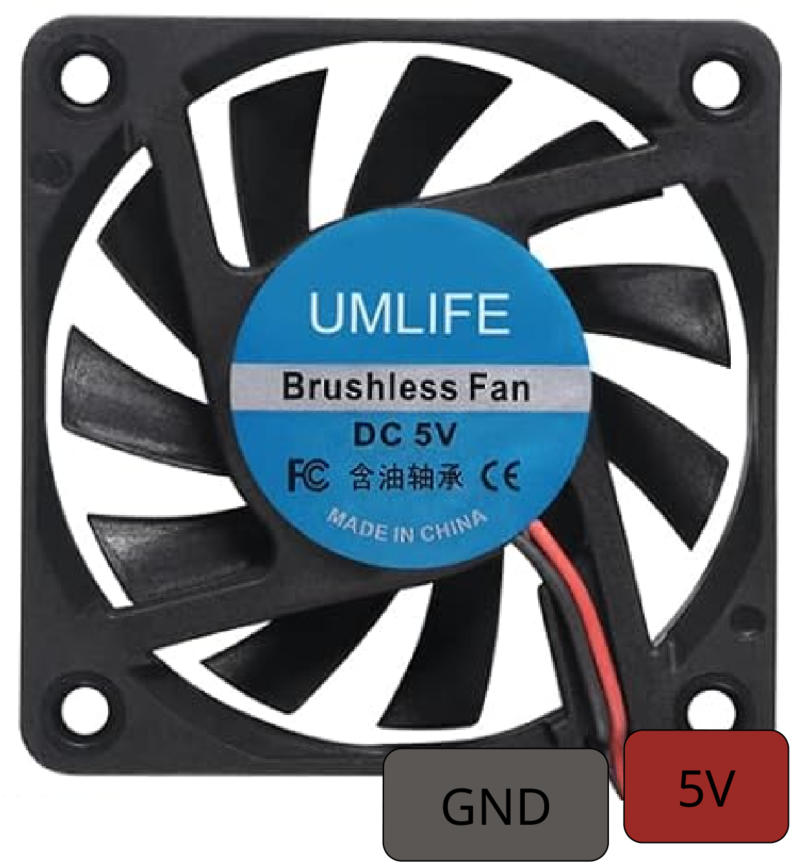
5 V brushless fan and its GND / 5 V cable labels.

**Figure 8.**
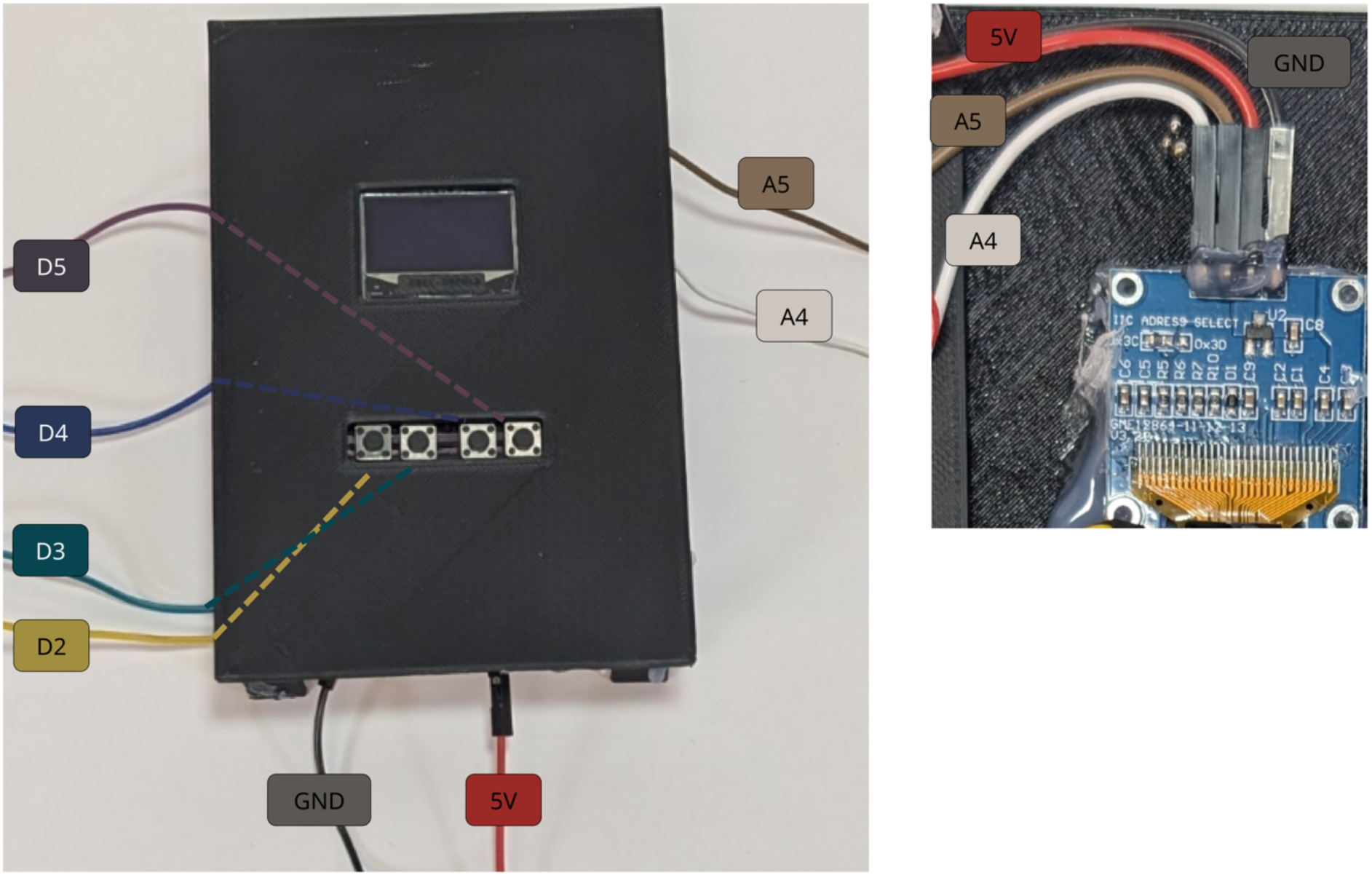
Combined front control/display panel and OLED-rear photograph, showing the control harness relationships. (Left) Annotated control/display panel (P8) with four button-signal leads and the OLED display supply lines. (Right) Reverse of control/display panel with close-up of the SSD1306 OLED connection: GND, VCC/5 V, SCL (A5) and SDA (A4).

**Figure 9.**
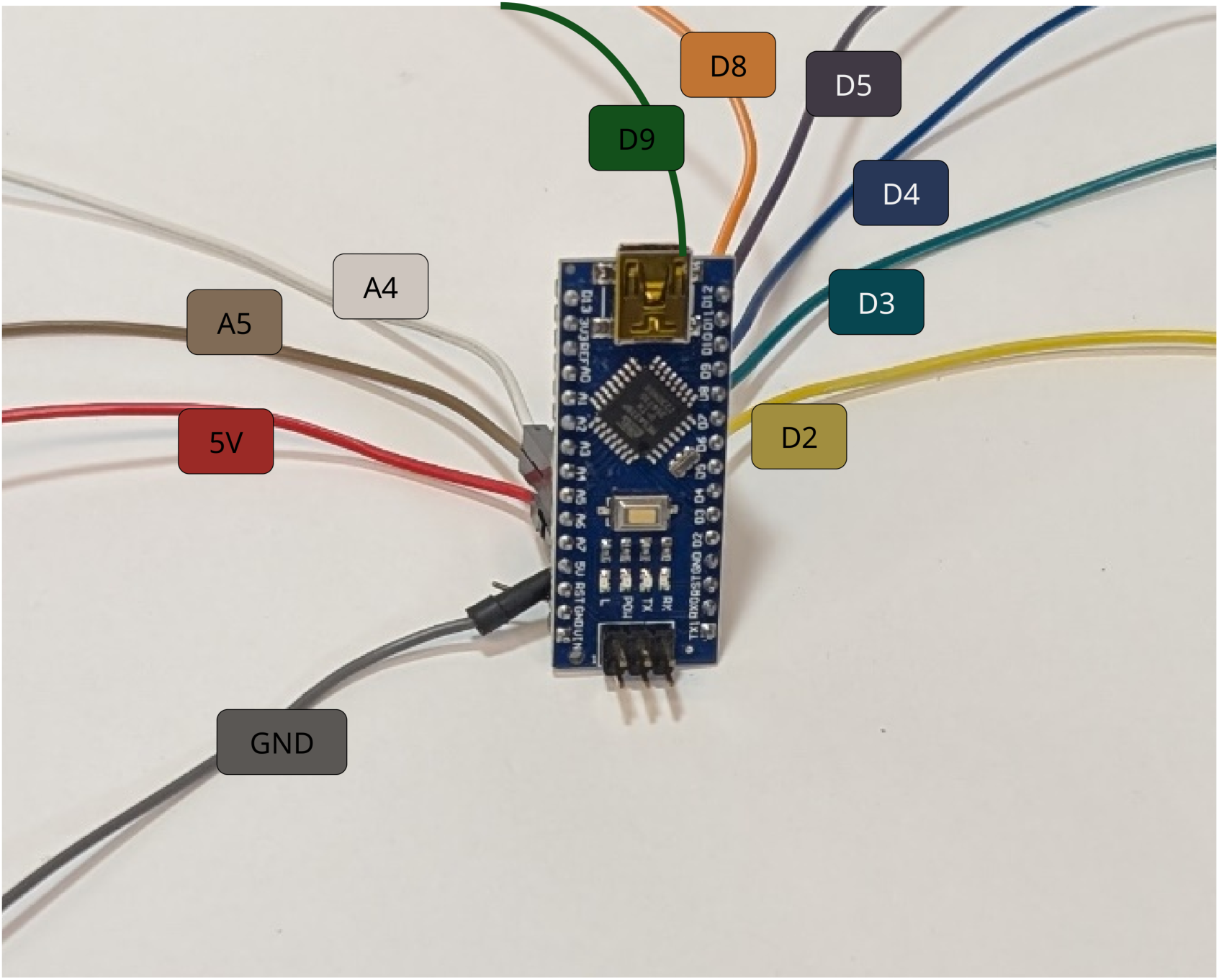
Annotated Nano-compatible controller showing D2–D5, D8/D9, A4/A5, 5 V and GND.

**Figure 10.**
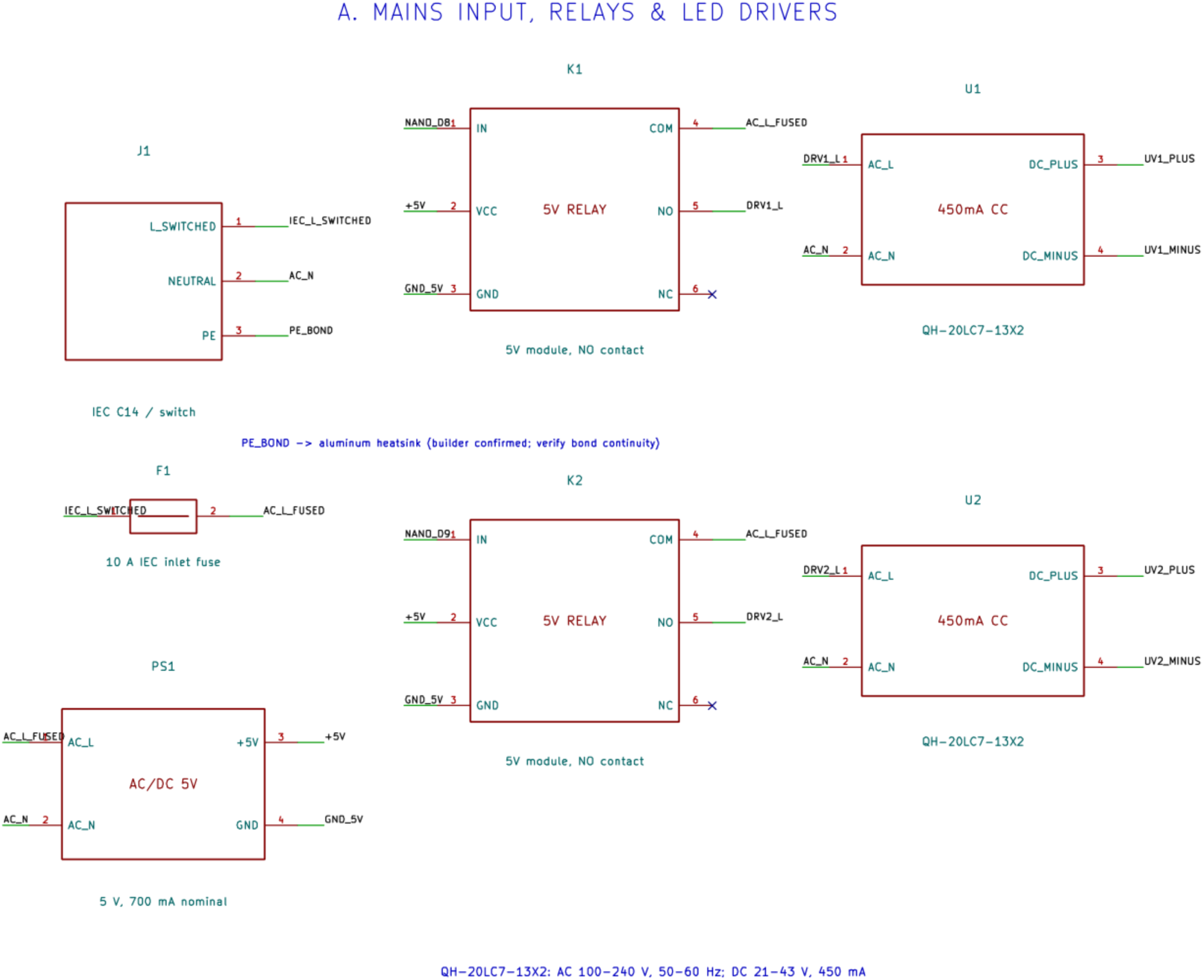
AC input, PE, fuse, two relay-switched driver inputs and 5 V conversion.

**Figure 11.**
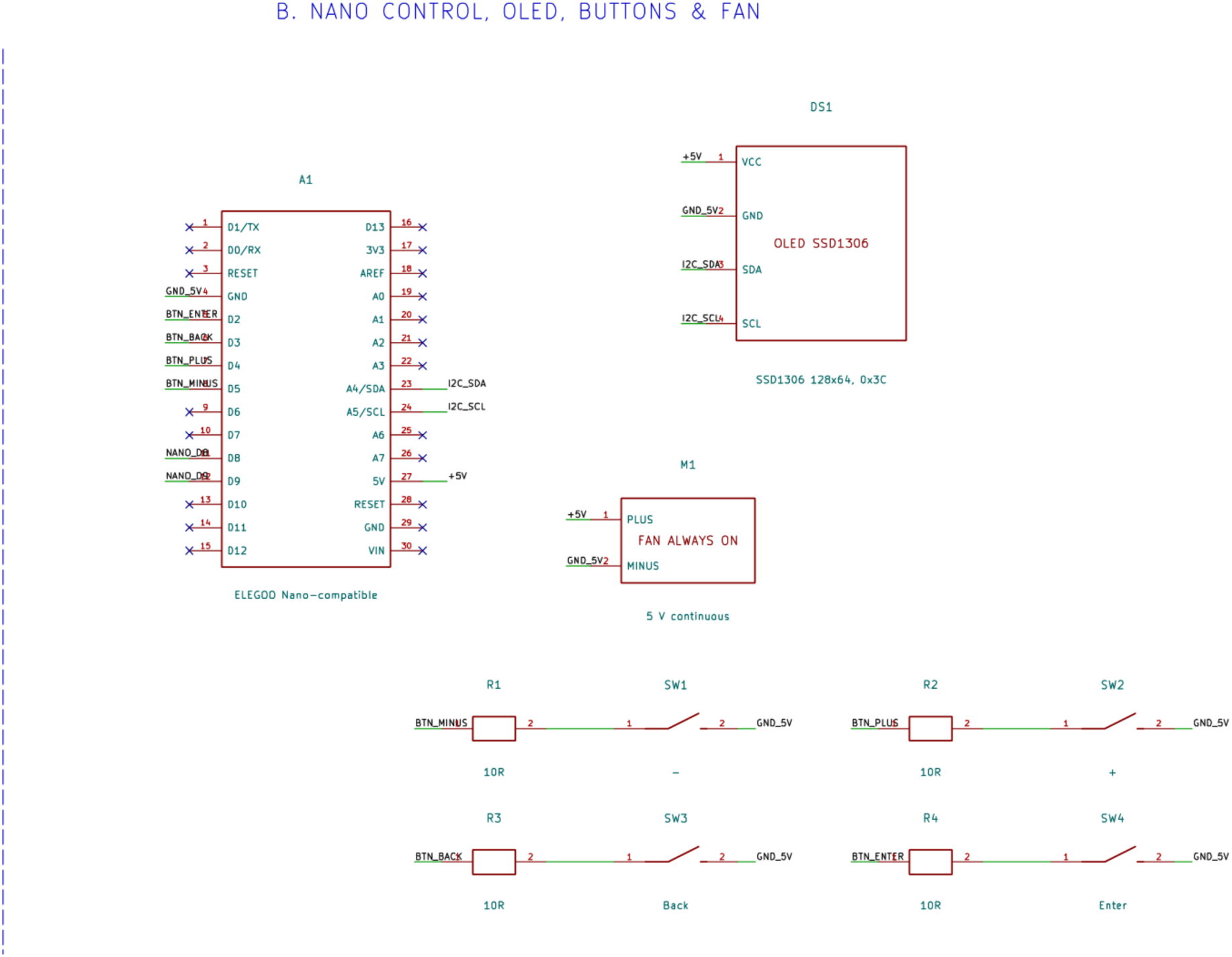
Nano pin functions, OLED I²C connection, four button-resistor paths and continuous fan supply.

**Figure 12.**
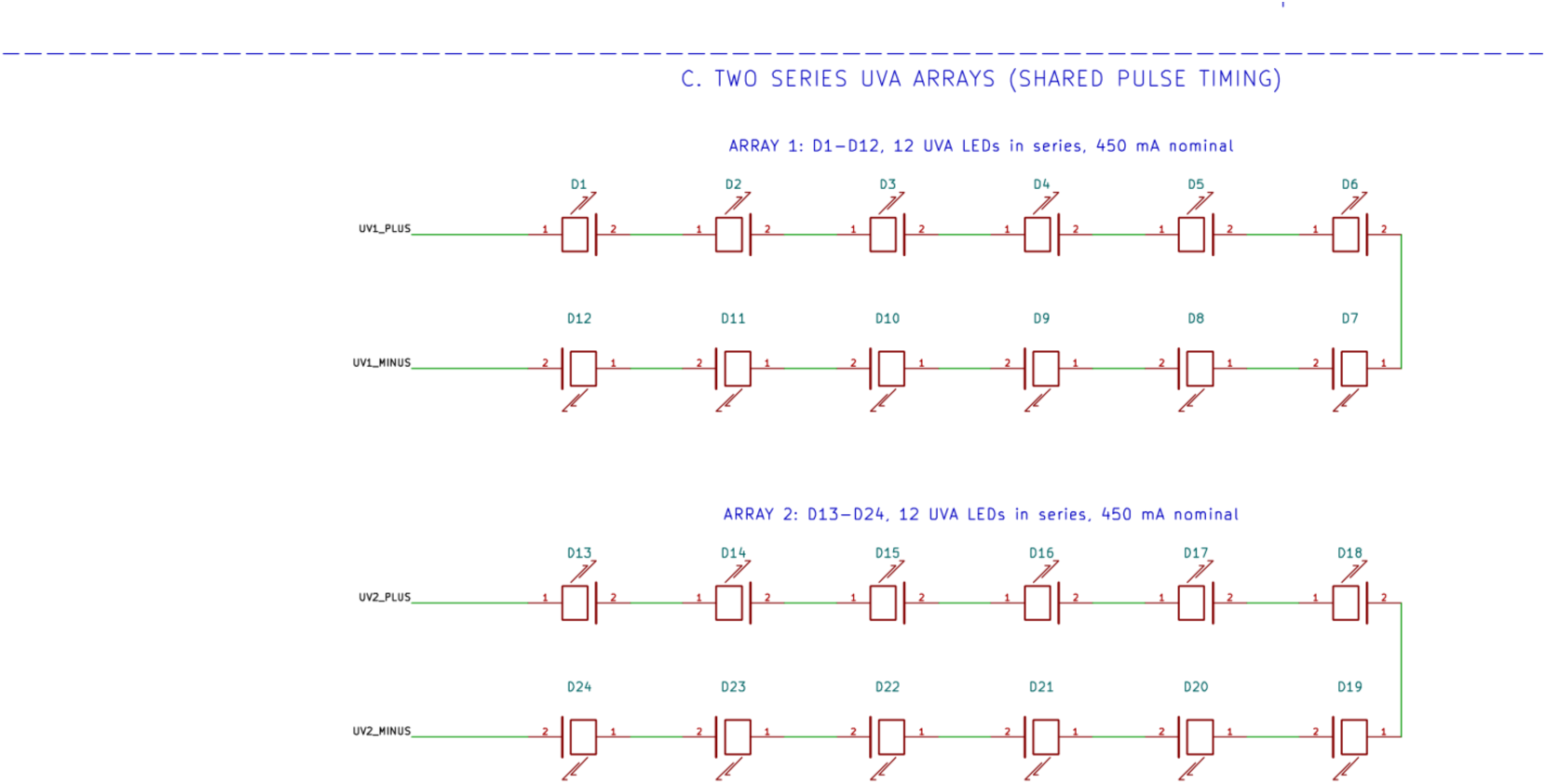
The two independent 12-LED series strings, one for each array.

**Figure 13.**
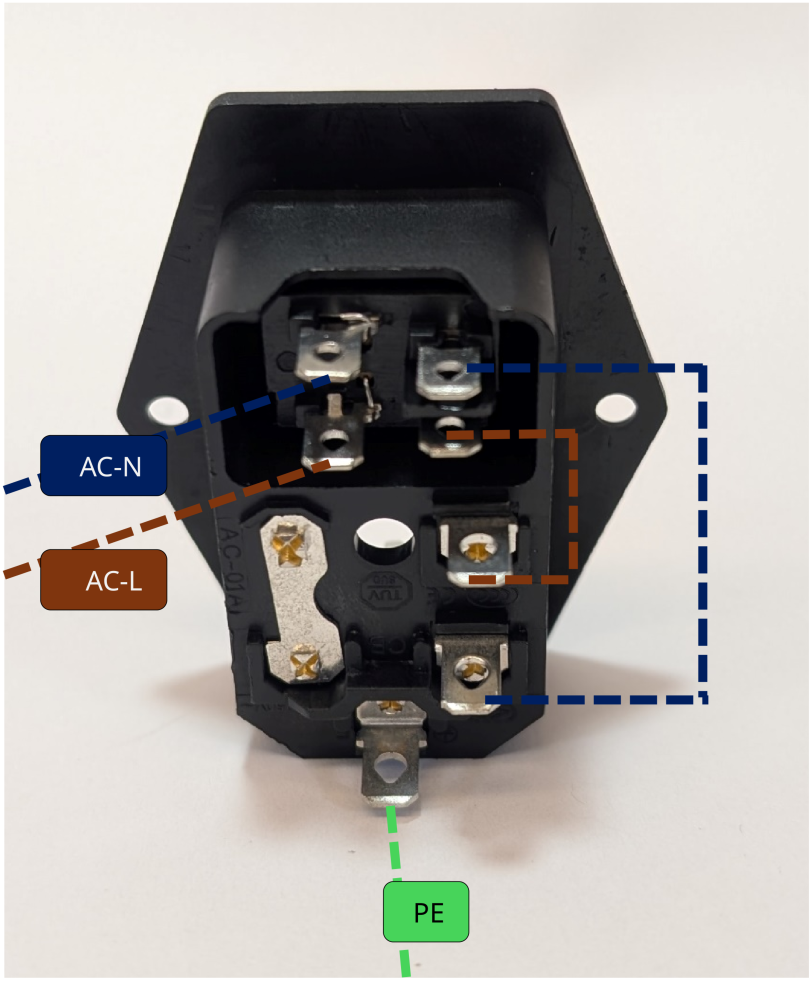
Rear of the switched IEC inlet with AC-N, AC-L and PE annotations.

**Figure 14.**
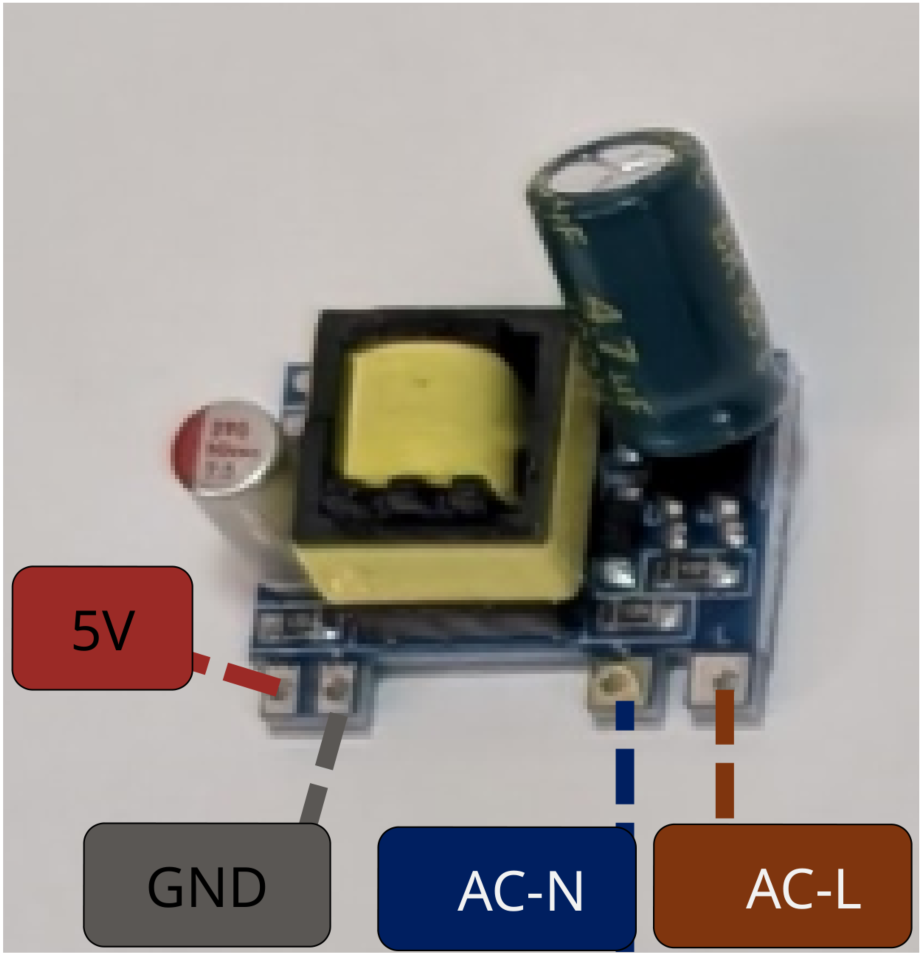
AC-input / regulated-5 V converter with input and output labels.

**Figure 15.**
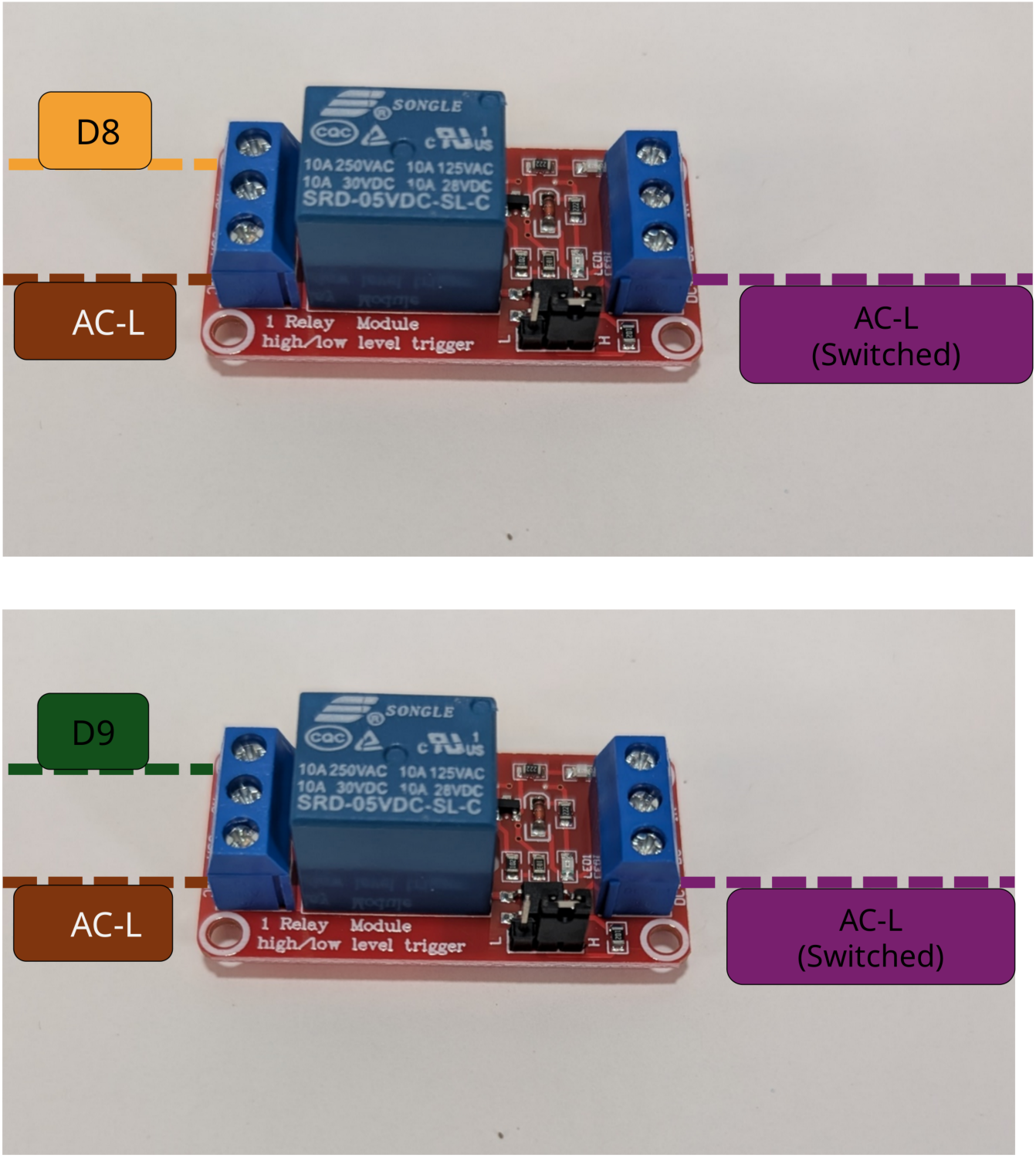
Top-array and bottom-array relays as annotated: D8 signal, incoming AC-L and switched AC-L out. The bottom-array relay is identical except that D8 signal is substituted by D9.

**Figure 16.**
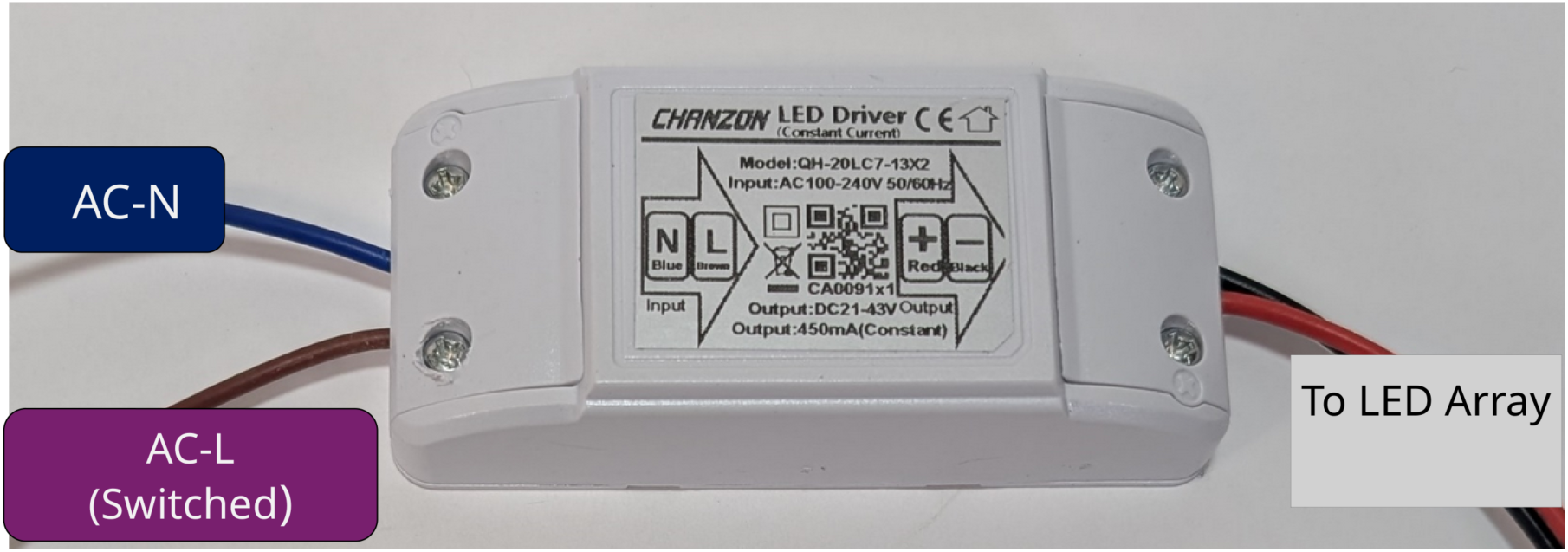
Constant-current driver: AC-N input, relay-switched AC-L input, and dedicated LED-array DC output.

**Figure 17.**
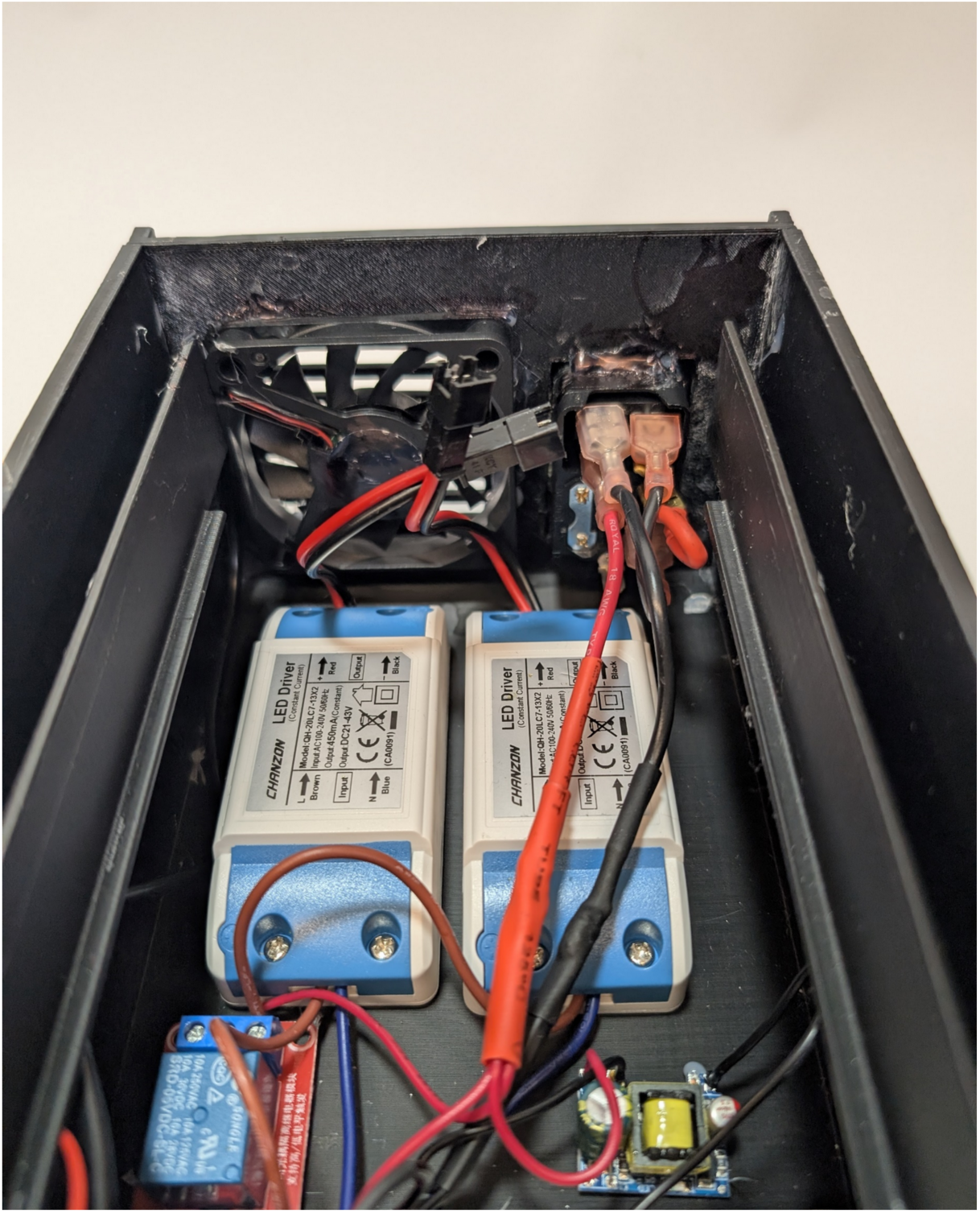
Lower Main enclosure interior showing fan, IEC inlet, two drivers, relay and supply placement. A second relay is placed below the one shown here.

#### 5.21 Operation

- With the device switched OFF, install the plate or tube carrier, confirm that the sample and carrier are seated, then move the cover to the half-closed stop that fully shields the UV/sample area.
- Power on the device using IEC inlet switch.
- Use Enter to access Active UV Arrays, then adjust Top/Bottom/Both, ON time, OFF time and Total Time with minus/plus and Enter. Use Back to return to the prior screen; read the confirmation screen before START.
- Press Enter to begin the run while the cover is HALF-CLOSED and the emitter region is already blocked. The selected array(s) enter the ON phase immediately. Slide the cover onward to FULLY CLOSED so the display is also covered.
- **DO NOT OPEN COVER WHILE LIGHT IS ON.** After the elapsed-time limit, make sure the outputs are OFF before uncovering the sample area. Pressing ‘Back’ during the active run commands a STOP.
- When using tubes or a tube insert, record the specific vessel/insert geometry and measure source-to-vessel distance; the 10 mm value applies only to the described 96-well sample-well bottom.

## 6. Operation instructions

### 6.1. Sample loading and irradiation

1. Place the device in its validated location, including inside a refrigerator at approximately 4 °C when using the low-temperature RNA protocol. Verify that the fan runs and that refrigerator airflow, device placement, and power feed routing satisfy laboratory electrical and UV-safety requirements.
2. Prepare the defined RNA/crosslinker samples in approved vessels, with matching dark and no-crosslinker controls. Record vessel, sample volume, optical path and sample position.
3. With the LEDs off, load the plate or tube insert and replace the sliding cover in the half-closed position, ensuring LED array is covered fully.
4. Use ENTER to advance through Active UV Arrays (Top, Bottom, Both), ON Time in seconds, OFF Time in seconds and Total Time in minutes, adjusting with the minus/plus keys; use BACK to return. Check the confirmation screen and press ENTER to start. Immediately slide cover to fully-closed position.
5. The run begins in the illuminated phase, follows the programmed on/off pattern and stops at the total elapsed-time limit. Pressing BACK during a run stops both arrays; record aborted runs separately.
6. Confirm the light source is off before opening the cover. Remove samples and proceed with the validated RNA cleanup and analysis.

### 6.2. Operating restrictions and protection

Verify enclosure is intact and grounding contacts are secure before connecting to power. Ensure sliding cover is in place before energizing UV arrays. Do not operate device with sliding cover removed. Take care to ensure device does not come into contact with spilled liquids or condensation drips in refrigerated environments.

## 7. Validation and characterization

### 7.1. Characterization of UV LED system

The QuickXL device utilizes commercially available, high-power UVA light-emitting diodes (Cleiscry, 3 W UV LED, B0CSSKPVHC) with a manufacturer-advertised emission wavelength of 365–370 nm. Each LED has a nominal maximum electrical power rating of 3 W, a recommended operating current of 450 mA, a forward voltage of 3.0–3.8 V, and a specified emission angle of 120–140°. The LEDs are mounted on 20 mm printed circuit boards and arranged into two independently powered and controlled arrays.

The LED arrays are operated using constant-current drivers and mounted to a common heat sink with continuous forced-air cooling to minimize thermal accumulation during irradiation. Samples are positioned approximately 10 mm from the LEDs. To accommodate temperature-sensitive RNA crosslinking experiments, the device can be operated within a 4 °C refrigerator. Under these conditions, continuous fan operation maintains the heat sink and sample plate near the ambient refrigerator temperature throughout a 10-minute irradiation, whereas operation without active airflow results in a steady temperature increase.

The optical emission spectrum of the LED arrays was experimentally characterized to determine their peak emission wavelength and spectral distribution, which were found to be λ_max_ = 371 nm, with a FWHM of 9.0 nm. A graph of the device output spectrum is provided in Figure 18. These measurements were used to characterize the UV exposure conditions independently of the manufacturer’s nominal electrical power specifications.

**Figure 18.**
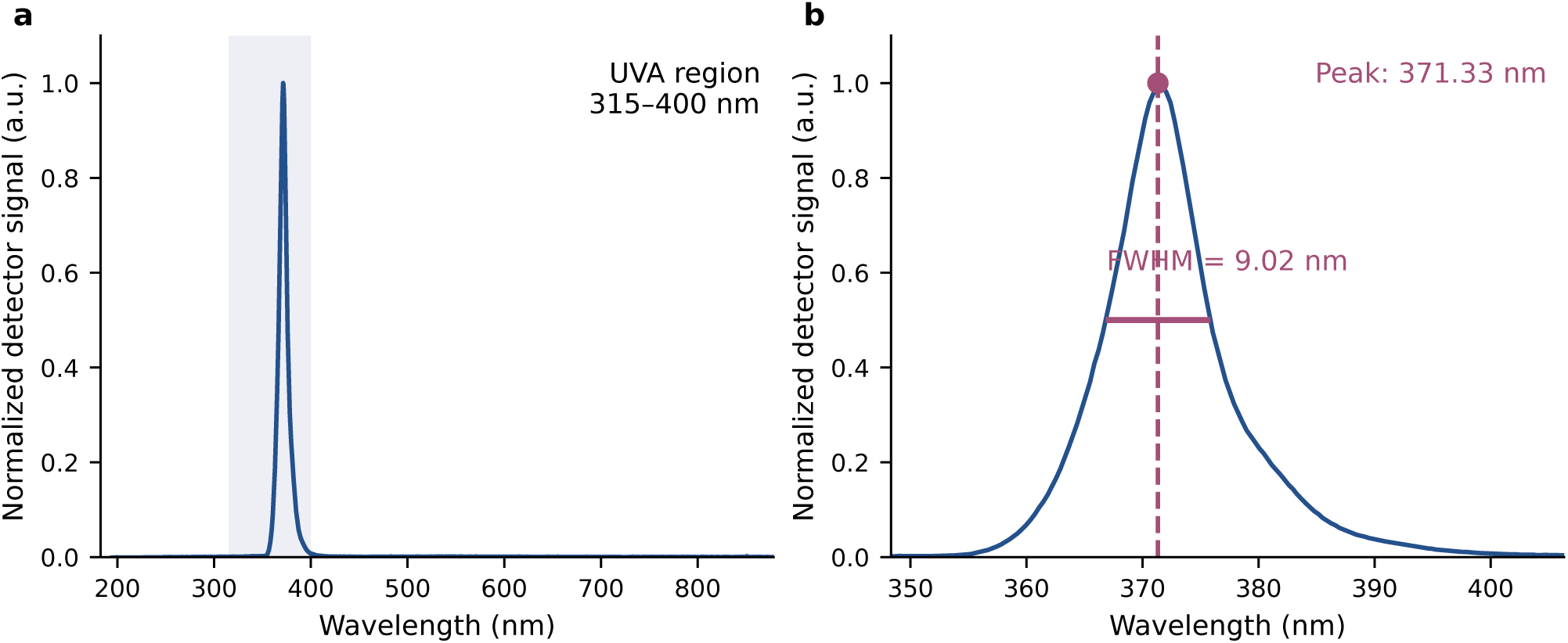
Spectral characterization of the QuickXL UVA irradiation device. (a) Measured emission spectrum of the UV LED source across the 200–850 nm wavelength range, with the UVA region (315–400 nm) shaded. The emission is concentrated in a narrow band near 370 nm, with approximately 96% of the integrated, background-corrected detector signal falling within the UVA region. (b) Enlarged view of the principal emission band showing a peak wavelength (λ_max_) of 371.33 nm and a full width at half maximum (FWHM) of 9.0 nm, corresponding to half-maximum wavelengths of 366.8 and 375.9 nm. Spectral measurements were background-corrected and normalized to the maximum recorded intensity.

### 7.2. Thermal performance

Maintaining stable sample temperatures during UV irradiation is particularly important for temperature-sensitive applications, including RNA crosslinking experiments, where heating may influence RNA structure or reaction kinetics. Although the QuickXL device employs high-power UVA LEDs, electrical energy not converted into light is dissipated as heat, potentially increasing sample temperatures during prolonged irradiation. In preliminary passively-cooled prototypes utilizing UVA LEDs, samples warmed notably across the treatment time; in some cases this effect was sufficient to produce sample vapor condensation on the plate lid. To minimize thermal accumulation, both LED arrays are mounted to a common aluminum heat sink with continuous forced-air cooling of the entire sample enclosure.

To evaluate the thermal performance of the device, sample temperature was monitored using a thermocouple immersed in 50 µL of water within a single well of a 96-well plate positioned on the irradiation platform. Temperature measurements were recorded at 1 s intervals during 10 min of uninterrupted LED operation under two environmental conditions: room temperature (approximately 20 °C) and refrigeration (4 °C). Refrigerated operation was evaluated because the device is designed to be placed directly inside a laboratory refrigerator for temperature-sensitive experiments, using the refrigerator to establish the ambient temperature while the integrated fan dissipates heat from the LED assemblies.

As shown in Figure 19, the device exhibited minimal sample heating under both conditions. During refrigerated operation, the sample temperature remained essentially constant, with an initial and final temperature of 4.2 °C and recorded values ranging from 4.1 to 4.3 °C throughout the irradiation period. The mean sample temperature was 4.2 °C, indicating that continuous LED operation did not produce appreciable heating above the refrigerated starting temperature.

**Figure 19.**
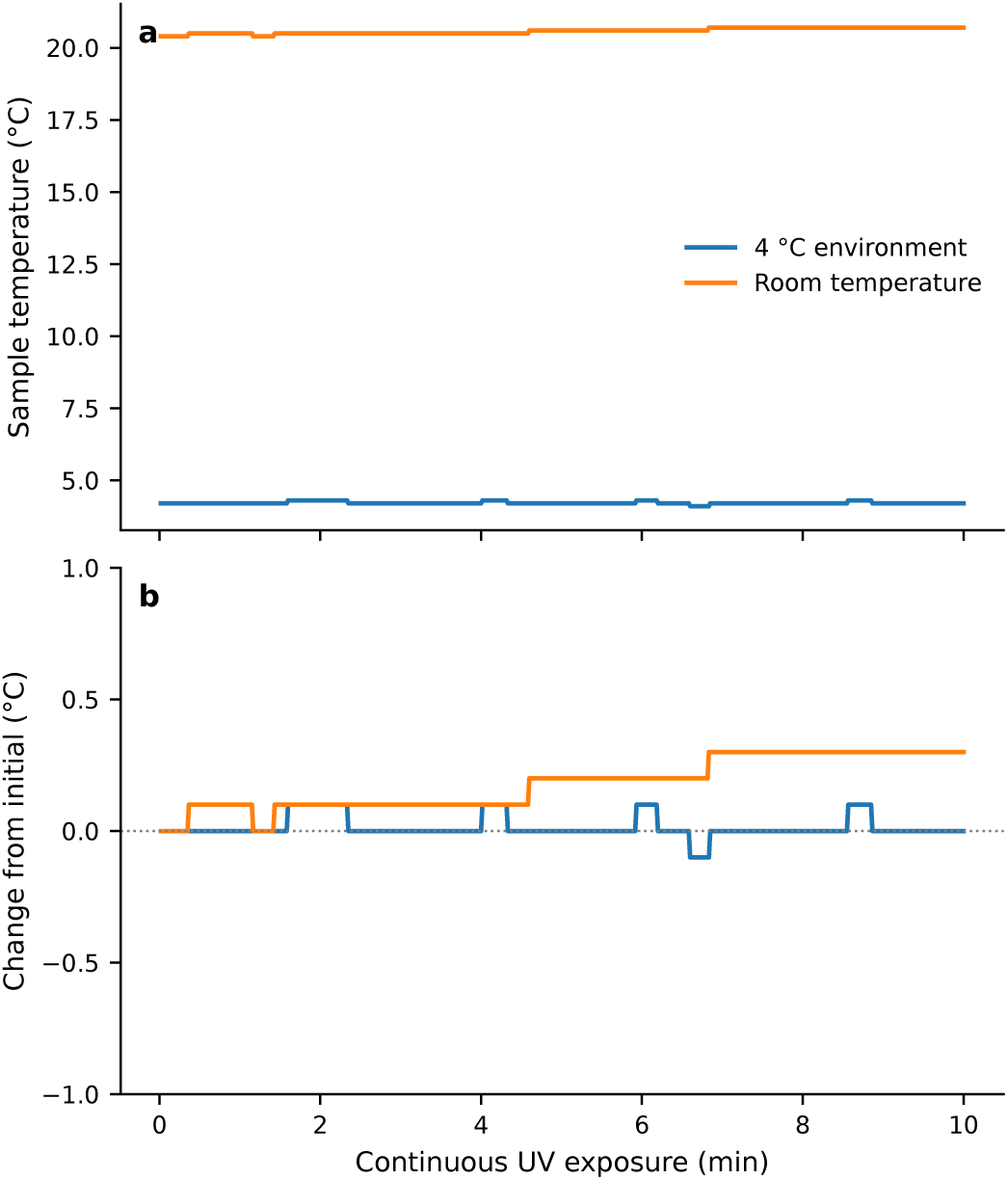
Thermal performance of the QuickXL UV irradiation device under refrigerated and ambient conditions. Temperature was measured using a thermocouple immersed in 50 µL of water in a single well of a 96-well plate positioned on the device during 10 min of continuous UV irradiation. Measurements were recorded at 1 s intervals with the device operated inside a 4 °C refrigerator or at room temperature (20 °C). **(a)** Absolute sample temperature during continuous irradiation. Under refrigerated conditions, the sample temperature remained near 4.2 °C, whereas the room-temperature sample increased gradually from 20.4 °C to 20.7 °C. **(b)** Temperature changes relative to the initial sample temperature for each condition. The refrigerated sample exhibited no net temperature increase, with fluctuations of approximately ±0.1 °C, while the room-temperature sample increased by 0.3 °C over the 10 min exposure. These results demonstrate minimal sample heating during continuous UV irradiation.

At room temperature, the sample temperature increased gradually from 20.4 °C to 20.7 °C over the 10 min exposure, corresponding to a net increase of only 0.3 °C. The mean temperature over the measurement period was 20.6 °C. This modest increase suggests that the combination of passive heat dissipation through the aluminum heat sink and continuous forced-air cooling is sufficient to limit sample heating during continuous irradiation under the tested conditions.

These results demonstrate that QuickXL can deliver sustained UV irradiation with minimal thermal disturbance to aqueous samples. In particular, operation within a standard laboratory refrigerator provides a straightforward means of maintaining samples near 4 °C without incorporating an active refrigeration system into the device itself. The low observed temperature increase also reduces the potential for thermal effects to confound UV-dependent experimental outcomes. Together, these characteristics support the suitability of QuickXL for applications requiring controlled UV exposure under stable temperature conditions.

### 7.3. Demonstration of psoralen-mediated RNA crosslinking

To demonstrate the utility of QuickXL for biological applications, the device was evaluated for psoralen-mediated RNA crosslinking and compared against two alternative UV irradiation systems: a handheld UV wand and a commercially available gel nail curing box. Psoralen-mediated crosslinking provides a useful model for assessing UVA irradiation performance, as psoralen intercalates into double-stranded nucleic acid regions and undergoes photochemical reactions upon UVA exposure, resulting in covalent crosslinks between complementary RNA strands. Crosslinking reactions were prepared using two complementary RNA fragments derived from the SARS-CoV-2 genome: a 42-nucleotide ORF1a fragment and a 36-nucleotide SL2/3 fragment. Each RNA oligonucleotide was present at a concentration of 100 ng/µL in LowTE buffer, with psoralen added to a final concentration of 0.2 mg/mL. Control reactions were prepared identically but without psoralen.

Following UV irradiation, reaction products were separated by denaturing polyacrylamide gel electrophoresis (PAGE) containing 8 M urea. Under these denaturing conditions, noncovalent RNA duplexes dissociate, allowing the covalently crosslinked RNA complex to be distinguished from the individual oligonucleotides as a higher-molecular-weight band. The handheld UV wand and commercial gel nail box were operated continuously for 20 min, whereas QuickXL was operated for 10 min using alternating 10 s irradiation and 10 s rest intervals, corresponding to 5 min of cumulative UV exposure. Samples in QuickXL were positioned approximately 10 mm from the LED arrays. Thus, QuickXL necessitated only one-quarter of the cumulative irradiation time used for either comparison device. As shown in Figure 20, all three devices produced a higher-molecular-weight RNA complex in the presence of psoralen, consistent with covalent crosslink formation between the two RNA fragments. However, substantial differences in crosslinking efficiency were observed. The handheld UV wand produced a relatively weak crosslinked band, while the commercial gel nail box generated a more prominent product. QuickXL produced the strongest crosslinked band, corresponding to an approximately 4.3-fold improvement in crosslinked product formation compared with the commercial gel nail box, despite requiring only one-quarter of the cumulative irradiation time. No corresponding crosslinked complex was detected in the absence of psoralen, supporting the specificity of the observed product for psoralen-dependent photochemistry.The enhanced crosslinking performance of QuickXL is consistent with efficient delivery of UVA irradiation to the sample through its closely positioned LED arrays. Importantly, the increased crosslinked product yield was achieved using an intermittent irradiation cycle rather than continuous exposure, demonstrating that effective RNA crosslinking can be obtained with relatively short cumulative exposure times.

**Figure 20.**
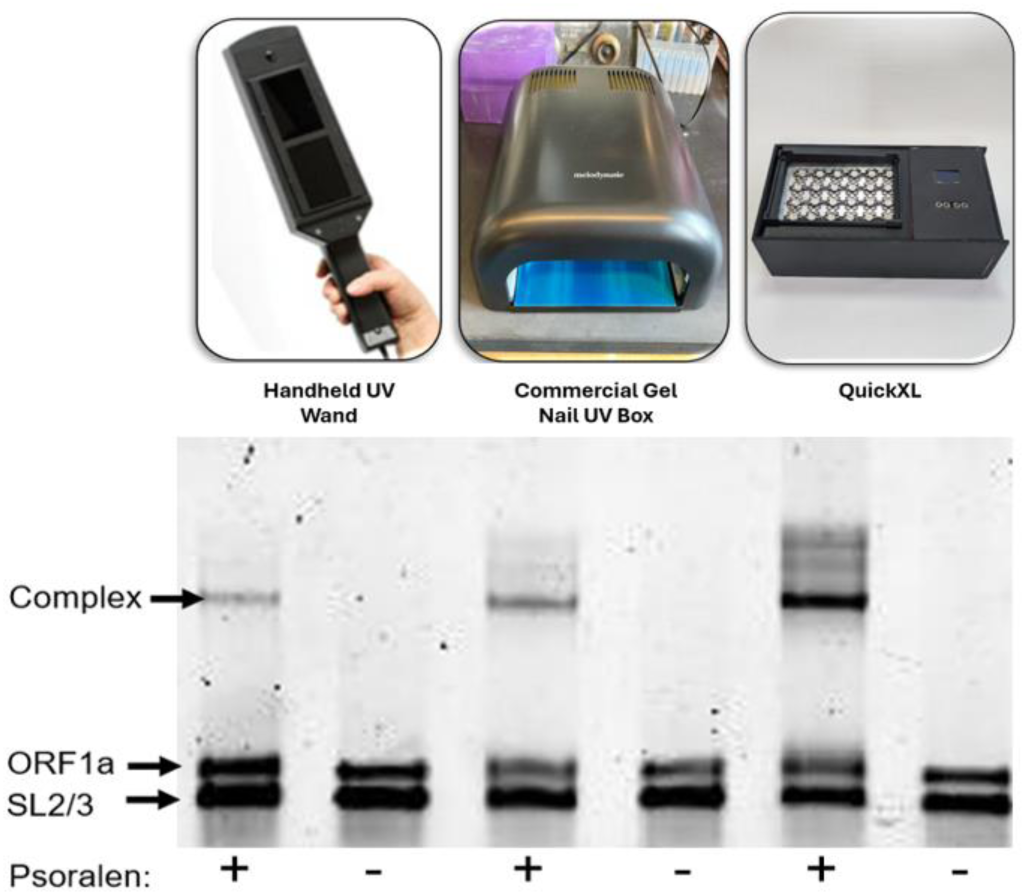
Comparison of psoralen-mediated RNA crosslinking efficiency using different UV irradiation devices. Representative 8 M urea denaturing PAGE analysis of RNA crosslinking reactions performed using a handheld UV wand, commercial gel nail UV box, or custom QuickXL irradiation device (photographs shown above the corresponding lanes). Reactions contained two complementary RNA oligonucleotides, ORF1a (42 nt) and SL2/3 (36 nt), each at 100 ng/µL in LowTE buffer, in the presence (+) or absence (−) of 0.2 mg/mL psoralen. The handheld UV wand and commercial nail box were operated continuously for 20 min, whereas QuickXL was operated for 10 min using alternating 10 s on/10 s off cycles, corresponding to 5 min of cumulative UV exposure. Bands corresponding to the individual RNA fragments and the higher-molecular-weight covalently crosslinked complex are indicated. QuickXL produced an approximately 4.3-fold increase in crosslinked product formation relative to the commercial nail box despite using one-quarter of the cumulative irradiation time. No corresponding crosslinked complex was detected in the absence of psoralen.

Together, these results establish the practical effectiveness of QuickXL for psoralen-mediated RNA crosslinking and demonstrate improved crosslinked product formation relative to the commercially available irradiation devices tested. The combination of efficient UVA delivery, programmable irradiation cycles, and temperature-controlled operation makes QuickXL particularly well suited for RNA structural studies and other applications requiring reproducible photochemical crosslinking under defined experimental conditions.

### 7.4. Conclusions

We have developed QuickXL, a low-cost, open-source UVA irradiation device designed to provide accessible, programmable, and reproducible photocrosslinking for nucleic acid research. The system combines 24 UVA LEDs arranged into two independently selectable arrays, microcontroller-based exposure control, a custom 3D-printed enclosure, and continuous heatsink cooling. The resulting instrument provides a versatile alternative to conventional UV irradiation systems, particularly for laboratories requiring controlled irradiation of multiple biological samples. By using commercially available electronic components and readily reproducible 3D-printed parts, QuickXL can be assembled and modified without specialized manufacturing equipment or custom printed circuit boards.

The primary application of QuickXL is psoralen-mediated RNA photocrosslinking, particularly for investigating RNA secondary and tertiary structure and interactions between distant regions of RNA molecules. Comparisons using a model RNA duplex system demonstrated substantially greater accumulation of psoralen-dependent crosslinked products with QuickXL than with a handheld UV wand or commercial gel nail UV box under the conditions tested. These results demonstrate the device’s suitability for RNA crosslinking experiments and suggest that its illumination geometry and operating characteristics can improve crosslink recovery relative to commonly available UV sources.

A particularly useful feature of QuickXL is its ability to operate in a refrigerated environment while maintaining continuous airflow across the LED heatsink. This arrangement provides a straightforward means of limiting sample heating during prolonged UVA exposure without the complexity or expense of an integrated thermoelectric cooling system. For RNA structural studies, maintaining samples near their intended experimental temperature is important because changes in temperature can influence RNA folding equilibria and potentially alter the interactions captured by photocrosslinking. The programmable illumination system also permits exposure to be divided into defined light-on and light-off intervals, providing flexibility in the temporal delivery of UV irradiation.

Beyond its immediate application to RNA structure analysis, QuickXL could be adapted for a variety of nucleic acid photocrosslinking workflows. Psoralen-based methods such as COMRADES, PARIS, and SPLASH rely on efficient stabilization of RNA duplexes prior to biochemical processing and sequencing [5–7], making controlled UVA irradiation an important component of these experiments. The device may also be useful for investigating DNA–DNA and DNA–RNA interactions, including studies of DNA interstrand crosslink formation, DNA damage and repair, and nucleic acid hybridization [12,13]. More broadly, programmable UVA exposure could facilitate smaller-scale investigations of intermolecular association and photocrosslinking kinetics. Beyond nucleic acids, the platform could be adapted for UVA-dependent photochemical applications, including photocleavage of light-sensitive chemical linkages and the modification of photoresponsive biomaterials [14]. Related photoinitiated reactions, such as thiol–ene photopolymerization, have also been employed in the fabrication of hydrogels and encapsulation of biologically active proteins [15]. Although these applications would require validation of wavelength compatibility, irradiance, and reaction conditions, QuickXL provides a potentially accessible platform for exploring such photochemical processes. Its compatibility with standard laboratory well plate sample formats may further facilitate parallel treatment of multiple experimental conditions, enabling systematic optimization of crosslinker or photoinitiator concentration, irradiation duration, exposure scheduling, and sample composition.

The modular construction of QuickXL also provides opportunities for further customization. Interchangeable sample holders could accommodate different microcentrifuge tubes, multiwell plates, or specialized reaction vessels, while alternative, easily-swappable LED configurations could provide different irradiation geometries, illuminated areas, or wavelengths. Such modifications would allow the system to be adapted to the requirements of individual experiments without extensive redesign of its electronic control architecture. The availability of editable enclosure models, electrical schematics, firmware, and assembly documentation is intended to support further development and adaptation by other laboratories.

Overall, QuickXL demonstrates a practical approach to constructing a customizable UVA irradiation platform using inexpensive, readily available components. Its combination of programmable exposure control, adaptable sample geometry, and passive thermal management makes it particularly suitable for psoralen-mediated nucleic acid crosslinking and related photochemical applications. By providing an openly documented alternative to commercial irradiation instruments, QuickXL has the potential to improve accessibility and experimental flexibility in RNA structural biology while supporting further development of reproducible, application-specific laboratory hardware.

## Ethics statements

No human or animal subjects were involved in the presented work.

## CRediT author statement

Ethan Kimmett: Conceptualization, Methodology, Software, Validation, Formal analysis, Investigation, Data curation, Visualization, Writing – original draft, Writing – review & editing

Victoria DeRose: Writing – review and editing, Resources, Funding acquisition, Project administration

## Acknowledgments

Funding: NSF (CHE2034277 and CHE 2109255 to VJD); University of Oregon Institute of Molecular Biology

